# Eudesmol-A promising inhibitor for glucosyltransferase: Docking and Molecular dynamics study

**DOI:** 10.1101/2021.12.27.474310

**Authors:** Shruthi N, Nithyashree R, Elakkiya Elumalai, Krishna Kant Gupta

**Affiliations:** School of chemical and Biotechnology, SASTRA Deemed University, India 613401; Centre for Bioinformatics, Pondicherry University, India 615014

**Keywords:** Tooth regeneration, infection, compounds, Eudesmol, Docking, Simulation

## Abstract

The loss of natural teeth can be avoided by invoking the molecular signal behind teeth regeneration. The destruction of the connective tissues is mainly due to bacterial origin which reacts to dental caries, a multifactorial disease. Glycosyl transferase is the enzyme which is involved in the glycosidic linkage. Glucosyltransferase inactivation reduces dental caries. This enzyme is a crucial virulence factor of *Streptococcus mutans*, a major pathogen that causes dental caries. In this present work, screening was done with library of anti-oxidant and anti-inflammatory molecules against the crystal structure of the target protein. Based on the predicted binding affinities, small molecules were selected and evaluated for their activity. Further, attempts were done to evaluate the toxicity of the lead compounds and compounds with no toxicity and good binding affinity were subjected for simulation and compared with reference complex. The potential energy of Glycosyl transferase-Eudesmol (proposed compound) (−1500 kj/mol) indicates its higher stability as compared to Glycosyl tranferase-G43 (reference) complex (−1100kj/mol).The inactives and actives compound for Glycosyl transferase was predicted from DeepScreening server.

## 1. Introduction

Dental caries is caused by the acids made by the bacteria which break down the teeth. *Streptococcus Mutans* is the major pathogen that causes this disease. *S.mutans* produces sticky glycosyl glucan polymers, which facilitate the attachment of the bacteria to the tooth surface. The glucans are a major source of the biofilm matrix that shields the microbial community from oxidative and mechanical stresses. ^[1]^ Prevention of dental caries includes regular cleaning of the teeth and should take die low sugar and small amounts of fluorides. Across the world, approximately 3.6 billion people have this problem in their permanent teeth. ^[2]^

Tooth regeneration is one of the main goals of restoring the loss of natural teeth. The therapeutic strategies for tooth regeneration have moved to the cell-based approaches. Teeth are ectodermal organs derived from sequential interactions between the oral epithelial cells and cranial neural crest-derived mesenchymal cells. ^[3]^ Teeth is a unique organ that composed of both hard and soft tissues. For the regeneration of the entire tooth, we are in grained to believe that stem cells or other cells must be transplanted. Regeneration of teeth can be done by various ways. They are cell transplantation, cell homing etc [4].

Glucosyltransferase inactivation reduces dental caries [5]. Dental caries is the most common problem that are faced by the People of our county and many loss their teeth and go for artificial teeth. This problem made us to think about the regeneration of teeth by natural small molecules which may not have much side effects during the process of regeneration.

To screen the anti-oxidant and anti-inflammatory molecules against the target protein and predict the toxicity of the screened molecules with high binding affinity and perform simulation studies for the reference and docked complex.

## 2. Methodology

### 2.1 Target selection

The crystal structure of the target was obtained from Protein Data Bank with the PDB ID: **3AIC**.The target protein of the enzyme Glycosyl transferase which is the main target that causes dental caries.

### 2.2 Library of small molecules

Aroma DB which covers basic and advance information of aroma molecules and provides a platform for comparative analysis and quantitative structure aroma relationship studies (QSAR). The set of anti-oxidant and anti-inflammatory molecules were selected from various different natural resources for screening. Around 500 molecules of both anti-oxidant and anti-inflammatory was selected for screening. ^[6]^

### 2.3 Molecular Docking studies

The molecular docking was carried out using PyRx tool (https://pyrx.sourceforge.io/). The small molecules are screened against the target protein of the enzyme Glycosyl transferase. The grid box was set as X=25, Y=25, Z=25.The binding residues (Table 1)were found using the Cast P tool (http://sts.bioe.uic.edu/castp/index.html?3igg).

**Table – 1.**
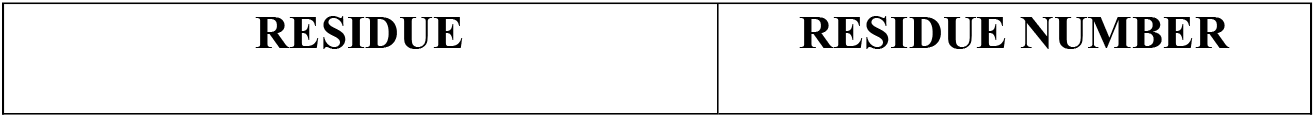
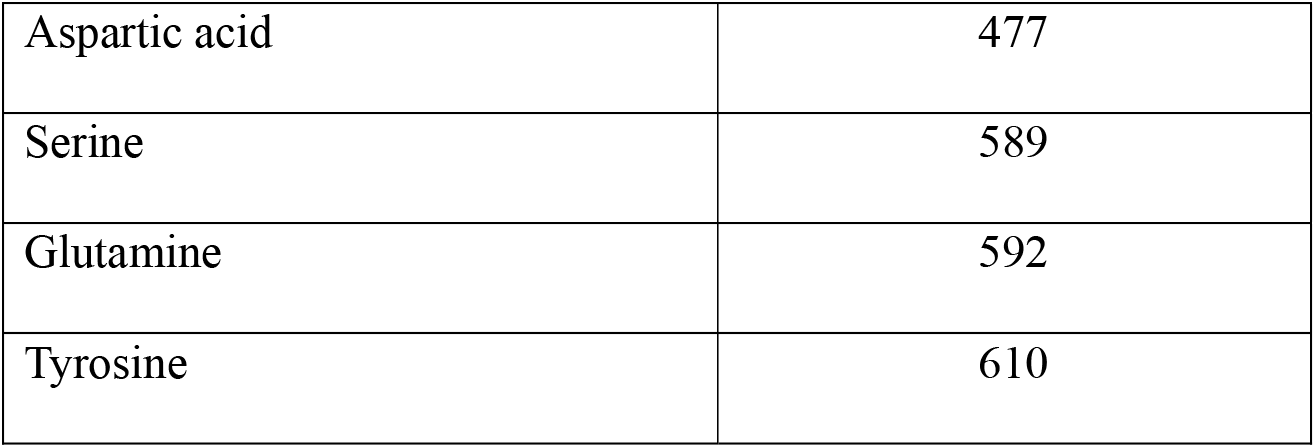
Binding sites of the protein.

### 2.4 Toxicity Estimation

The three compounds namely Beta Eudesmol, Beta Humelene and Beta Caryophyllene were identified with best binding affinities during the molecular docking studies. The above three compounds were used for the toxicity estimation using Tox Tree tool. Beta Eudesmol was having low toxicity levels when compared to the other two and thus it was taken for MD simulation [7].

### 2.5 Molecular Dynamic Simulation

The molecular stability of the 3-dimensional structure of the model protein was understood by performing the simulation using GROMACS v 4.5(https://www.gromacs.org/). The all – atom force field, GROMOS96 43a1 was implemented for the same. A periodic body with a dimension of 1.50nm was applied to the protein which was included in a cubic box. Sodium ions were used to neutralize the water molecules. Steepest descent algorithm and conjugate gradient method were used for the energy minimization of the system up to 50,000 steps. After minimization, equilibration of the systems were done for 100ps and restraining of the position done at the constant number of particles, volume and temperature i.e. NVT and also at constant number of particles, pressure and temperature ie. NPT. LINCS algorithm was used to constrain the bond angles and the water molecule geometry was used to constraint using SETTLE algorithm. The V – rescale weak coupling method was used to regulate the system temperature at 310K while Parrinello – Rahman method was used to set the pressure at 1 atm. A simulation run of 10ns along with 2fs time step was then set up with the system. Every 2ps structural co-ordinates were saved and the GROMACS package tools were used to analyze them. The protein structure with the minimum energy was retrieved finally.

MD simulation was carried out for Glycosyl transferase (target protein) for 10ns; Beta eudesmol (ligand) with protein for 4ns and G-43 (reference compound) with protein for 2.1ns.

### 2.6 Classification of small molecules using machine learning approaches

The inactives and actives compound for Glycosyl transferase were present in DeepScreening server [8]. We followed the following steps to build classification model.

**Selecting a model type:** Classification model was selected for the study.

**Compounds selection:** SDF file of small molecules (CHEMBL3988596) were selected with bioactivity information for Glycosyl transferase. There were Inactives 112 and Actives 392

**Feature generation:** There are 12 molecular fingerprints in deeplearning server. The “Klekota-Roth fingeprint (4860)” were selected for building classification model

**Klekota-Roth fingerprint:** SMARTS based substructure fingerprint based on Chemical substructures that enrich for biological activity. ^[9]^

**Hyper parameter selection: I**t was taken as by default. 30 cycles were there to improve the classification model.

**Submission:** Finally, it was submitted and classification model was predicted with score from 0 to 1. The more score refers active compounds. In contrast, less score means inactive compounds. The metrics and statistics of model was reported.

## 3. Results And Discussion

### 3.1 Molecular Docking

Molecular docking was performed for the reference molecule and the binding affinity calculated was −8.9kcal/mol (Table 2). The natural compounds that have higher binding affinity than the reference was used for further studies. The binding affinities of the compounds are listed below:

**Table – 2.** Binding Affinities of compounds.

| S.No | Ligand name | Target | Binding affinity (kcal/mol) |
| --- | --- | --- | --- |
| 1 | Beta-Eudesmol | 3AIE | -9.9 |
| 2 | Beta-Humelene | 3AIE | -9.4 |
| 3 | Beta-Caryophyllene | 3AIE | -9 |
| 4 | G43(Reference compound) | 3AIE | -8.9 |

### 3.2 Molecular Dynamic Simulation

#### 3.2.1. Molecular Dynamic Simulation of Protein

The potential energy of the protein was −1259.43 kj/mol. The protein is stable throughout the simulation (Fig. 4) and the binding affinity is highly negative.

**Figure – 1.**
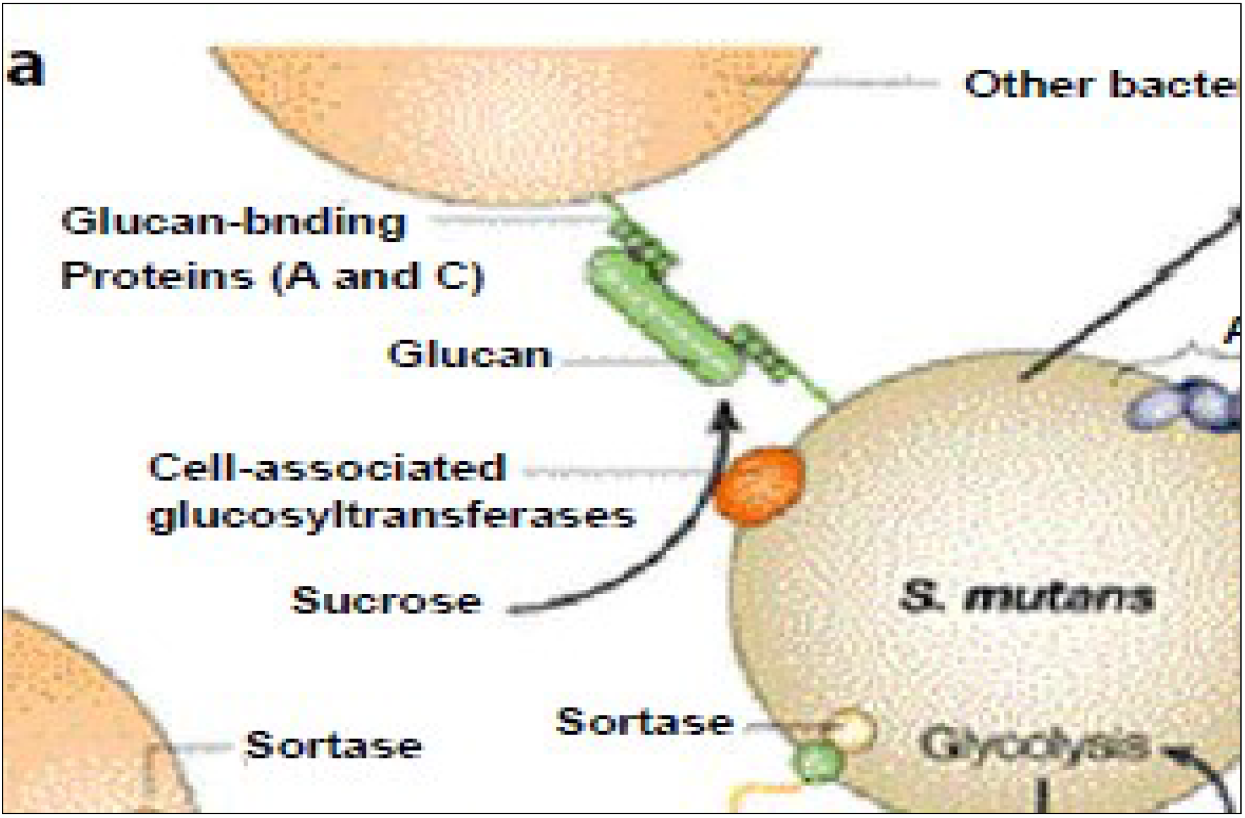
Role of *S.mutans* in dental caries.

**Figure – 2.**
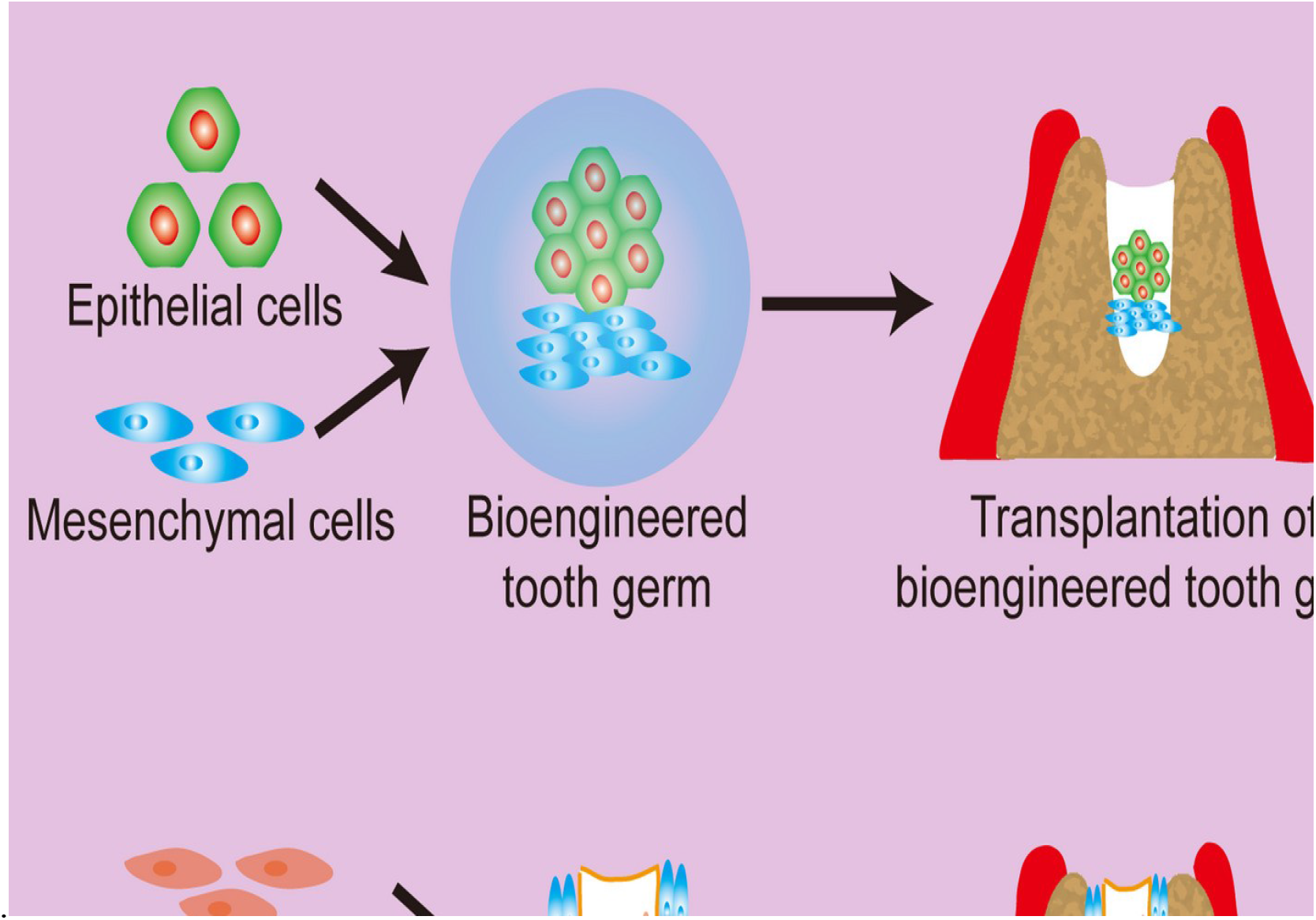
Stem cell-based tooth regeneration.

**Figure – 3.**
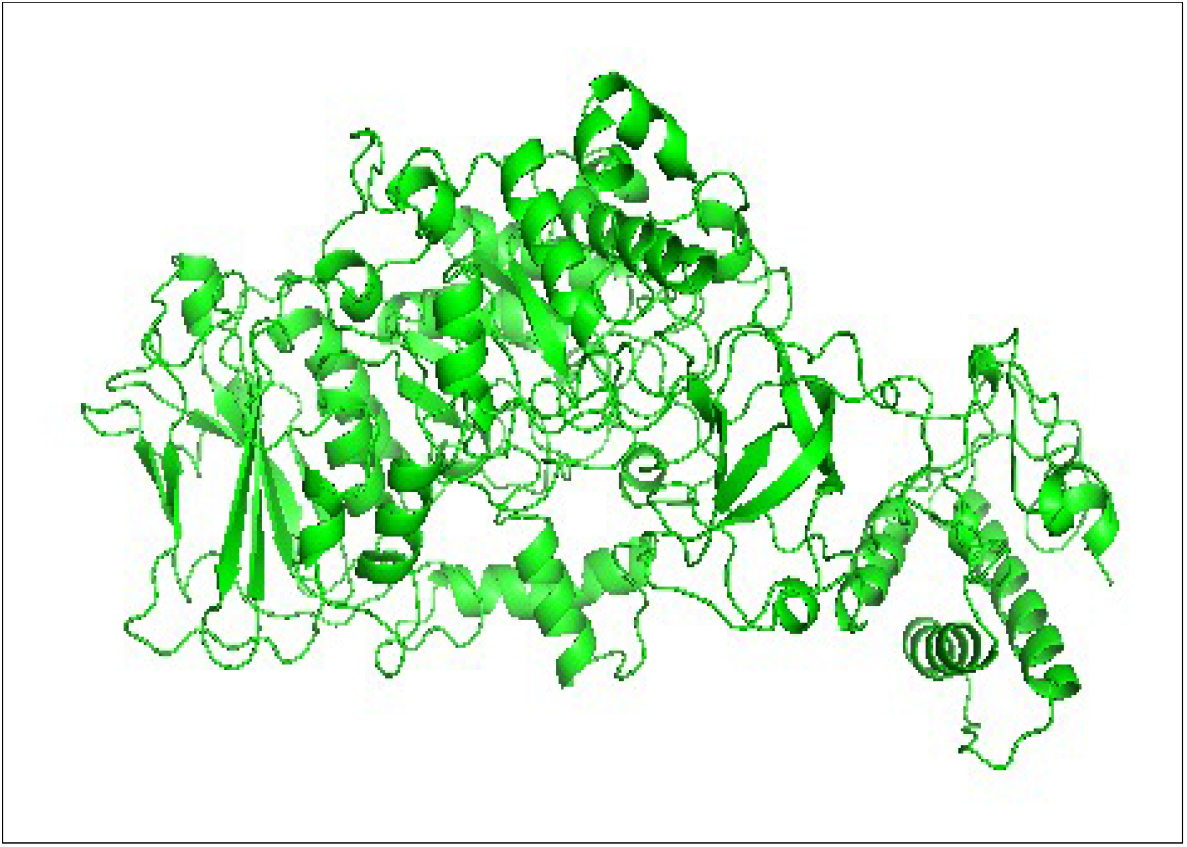
Crystal Structure of 3AIC.

**Figure – 4.**
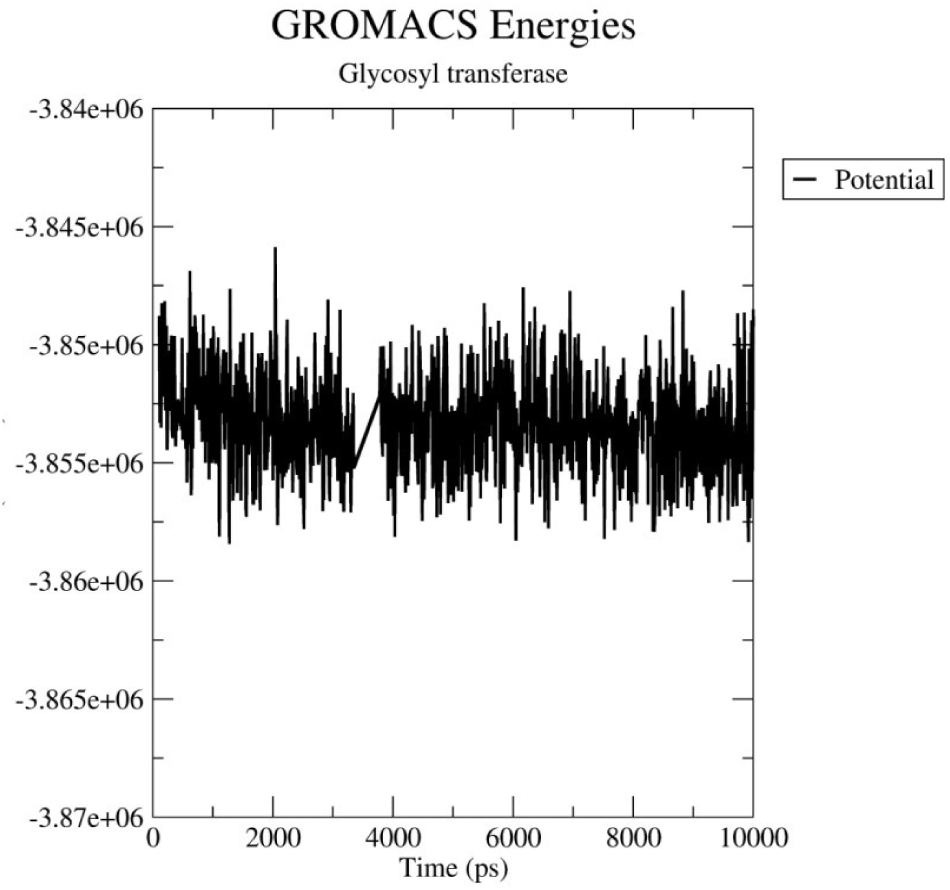
Potential energy graph of protein.

**Figure – 5.**
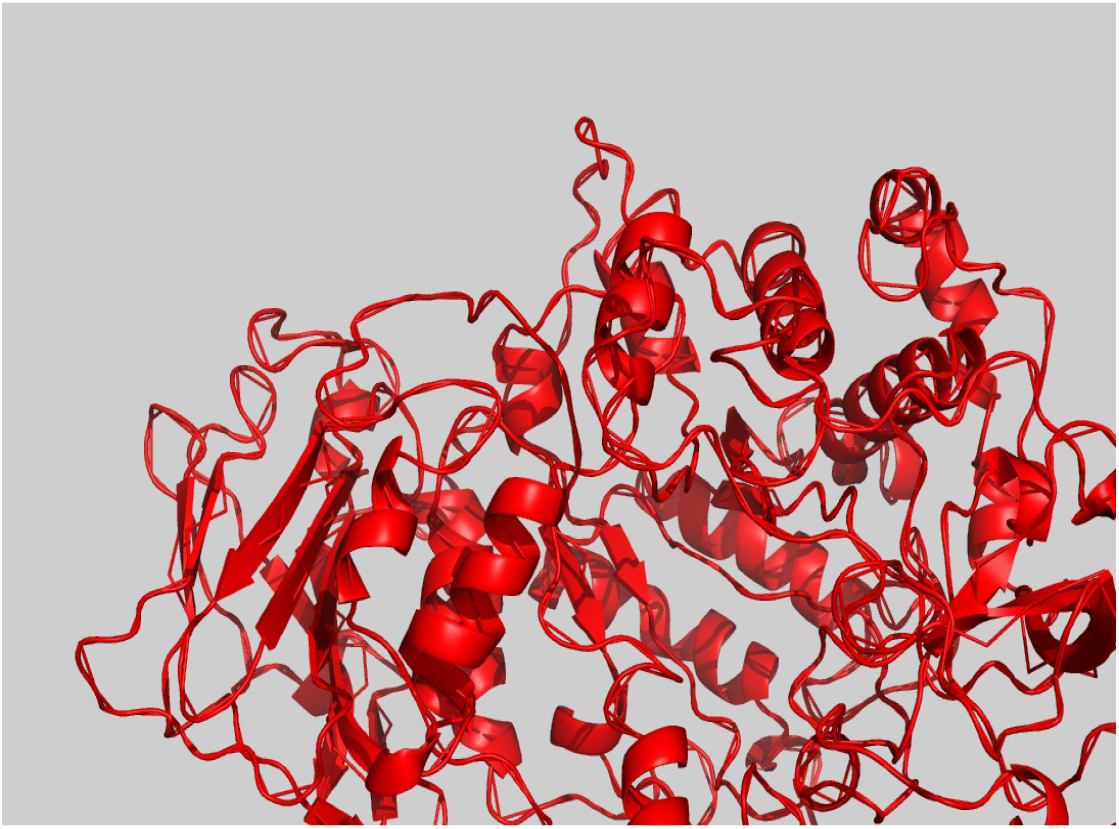
Lowest energy protein was attained at 2ns.

##### Kinetic energy, Enthalpy and Coulomb Short-Range energies

Kinetic energy, Enthalpy and Coulomb Short-Range energies are stable during the simulation process (Fig 6). The kinetic energy is positive because motion should always be positive since it’s a scalar quantity.

**Figure – 6.**
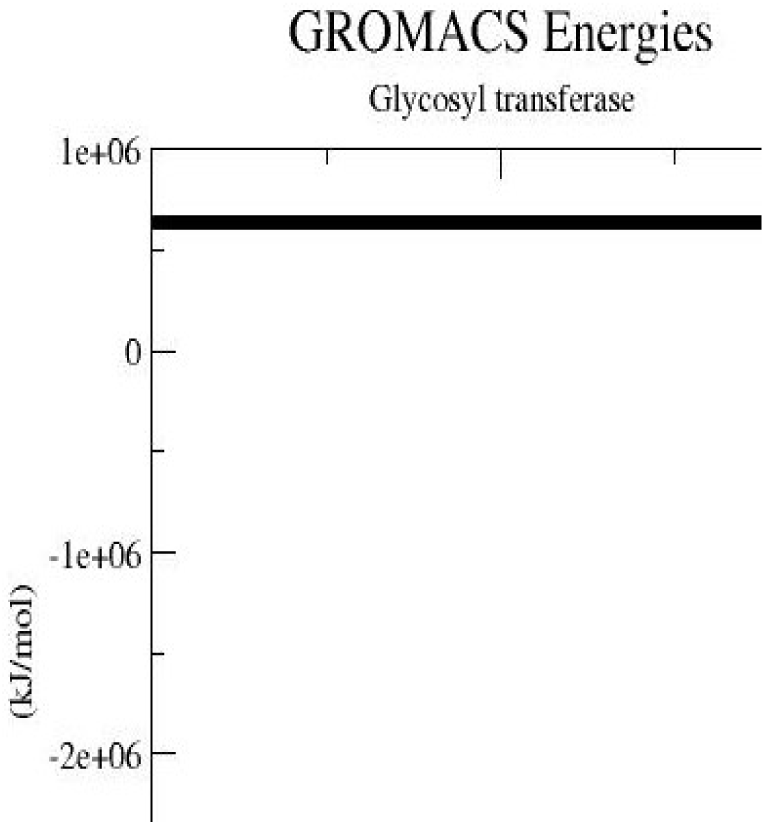
Energy graph of protein.

##### RMSD of Protein

The backbone RMSD profile of the protein shows that it reaches its equilibrium at 4-8 ns (Fig 7). There is no much fluctuation and hence its stable throughout the simulation.

**Figure – 7.**
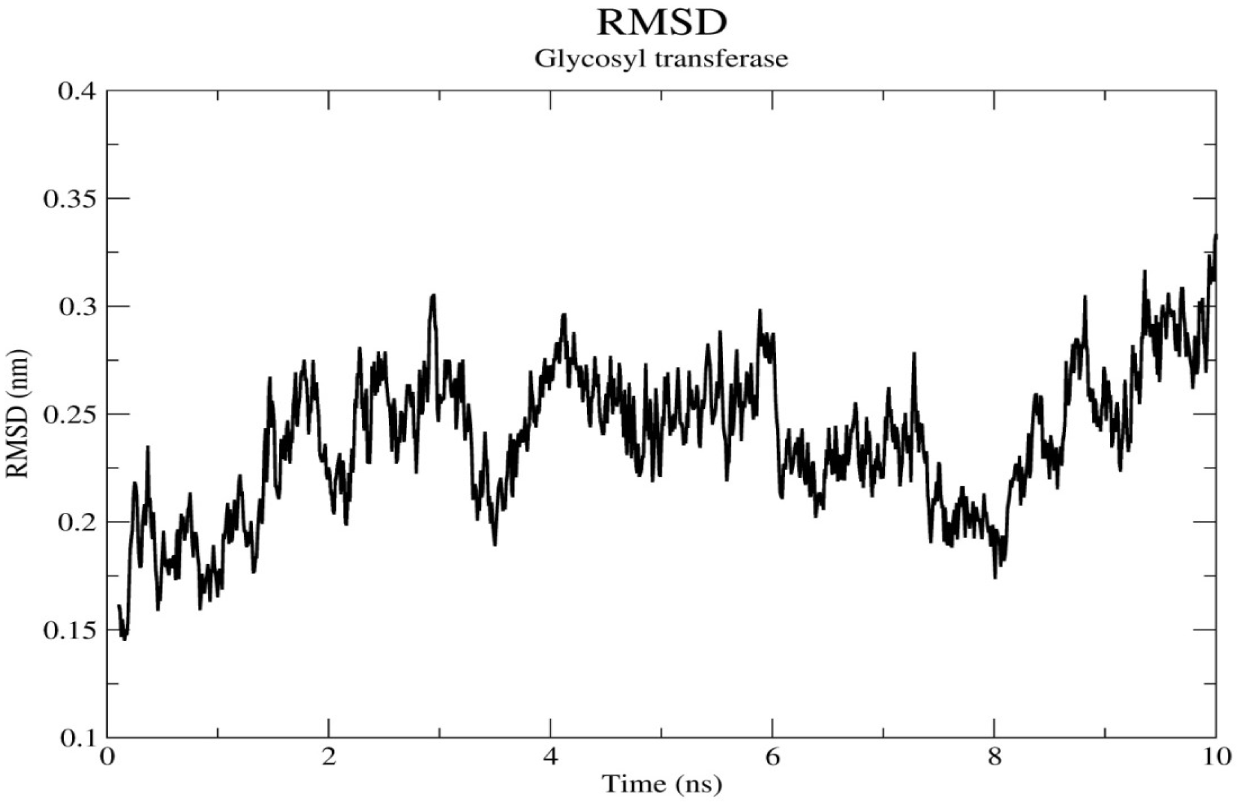
RMSD graph of protein.

##### RMSF of Protein

The RMSF profile of the protein was also evaluated to account for the flexible regions within the structure (Fig 8). The fluctuations are not seen much and hence it is stable.

**Figure – 8.**
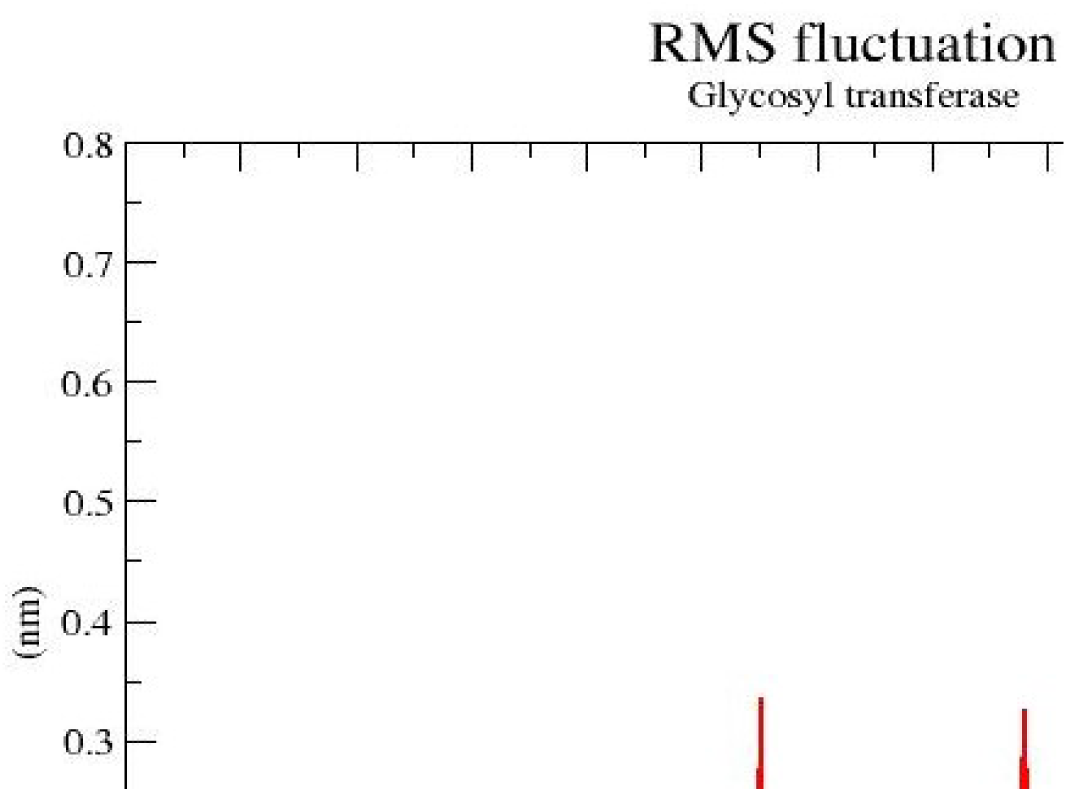
RMSF graph of protein.

##### Radius of Gyration of Protein

From the radius of gyration we find the compactness of the structure and the folding that was maintained during the simulation (Fig 9).

**Figure – 9.**
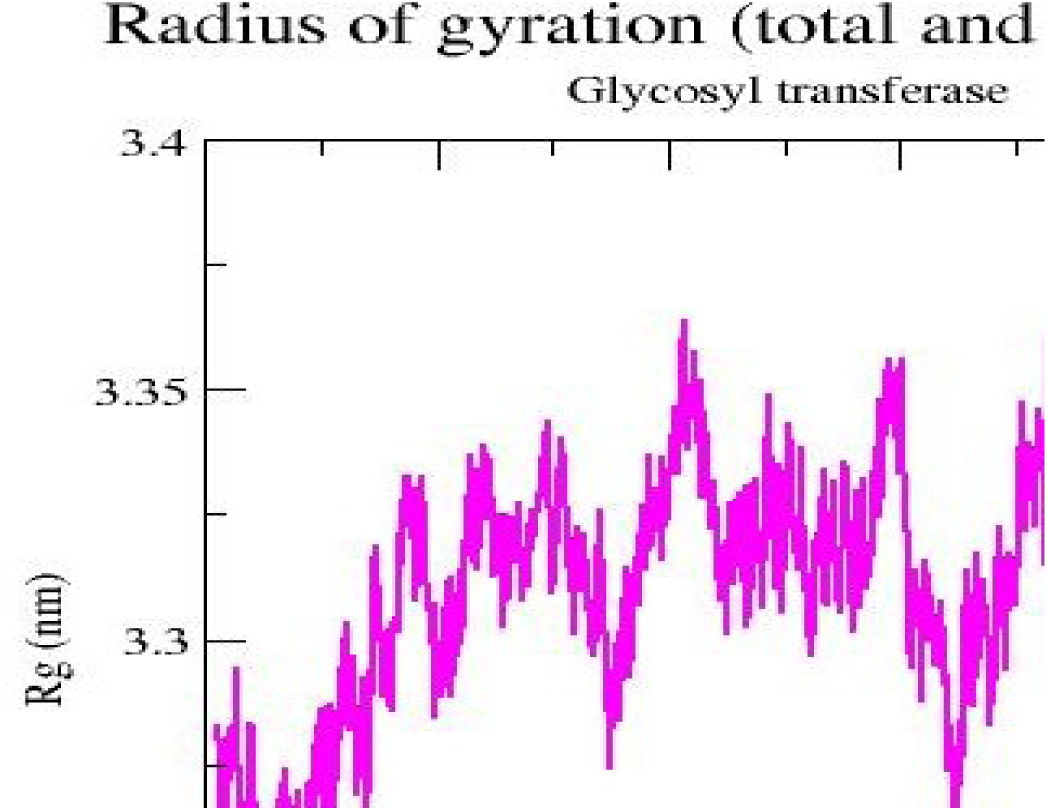
Radius of gyration graph of protein.

#### 3.2.2. Molecular Dynamic Simulation of Beta-eudesmol and protein

**Figure – 10.**
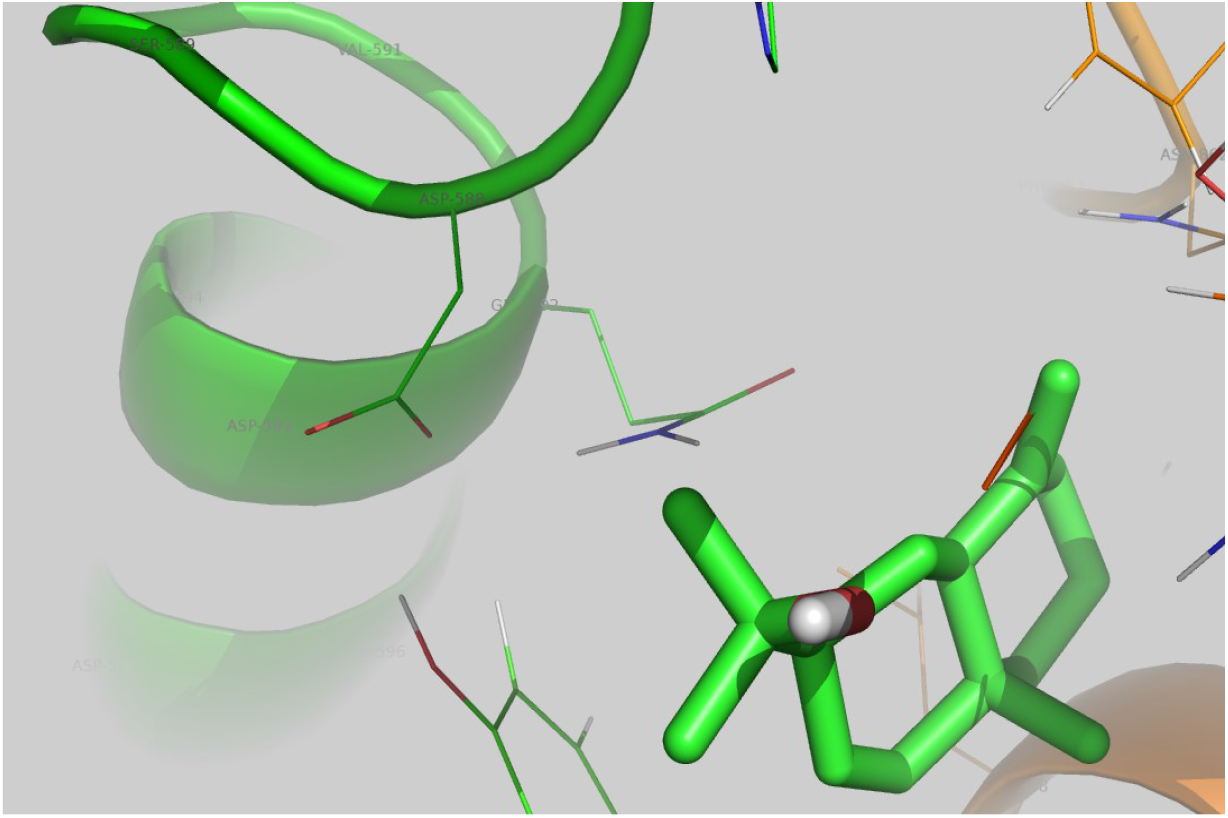
Structure of Beta-Eudesmol at 0ns.

##### Potential energy of Eudesmol

The potential energy of Eudesmol was found to be −1535.88 kj/mol (Fig 11).The potential energy of the complex is higher than the protein and hence the complex is more stable than the protein. The lowest energy was obtained at 3ns.

**Figure – 11.**
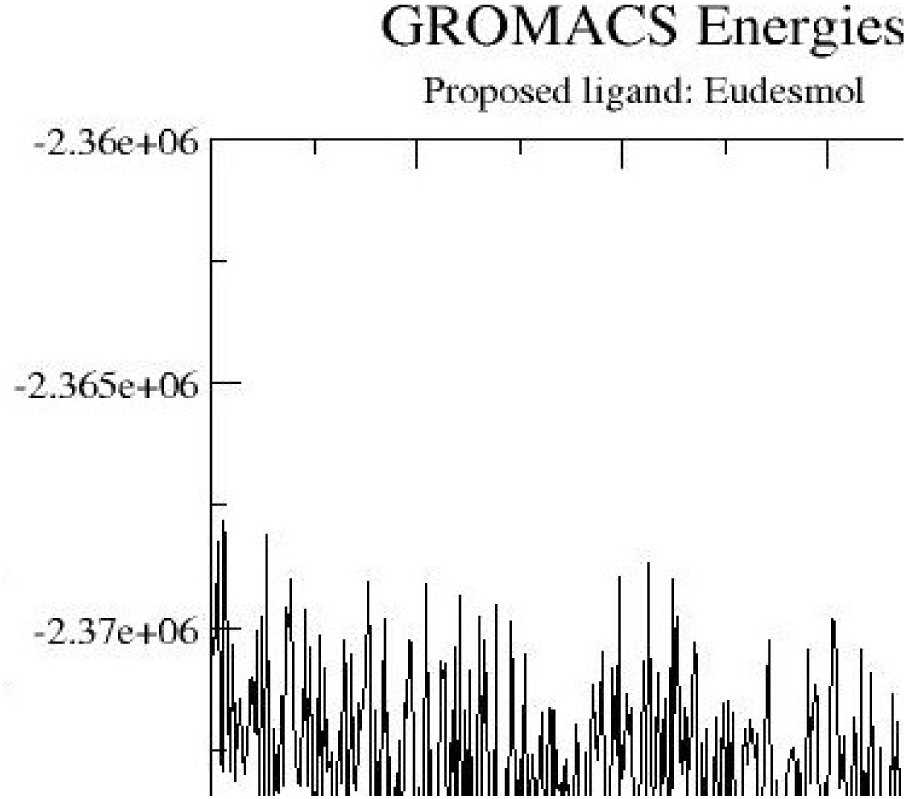
Potential energy graph of Beta Eudesmol.

##### Interaction energy of Beta Eudesmol

The number of H bonds is 2. From the graph it is possible to know about the interactions of the ligand with protein and the hydrogen bonds formed (Fig 12).

**Figure – 12.**
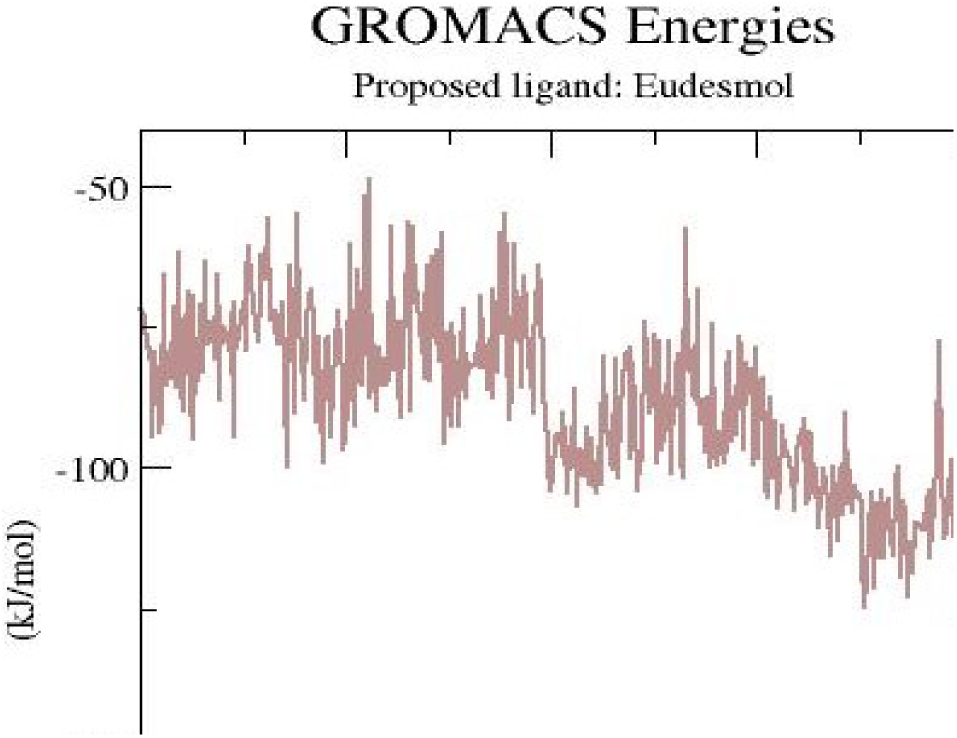
Interaction energy graph of Beta Eudesmol.

The interacting residues are **ASP588, GLN 592**

**Figure – 13.**
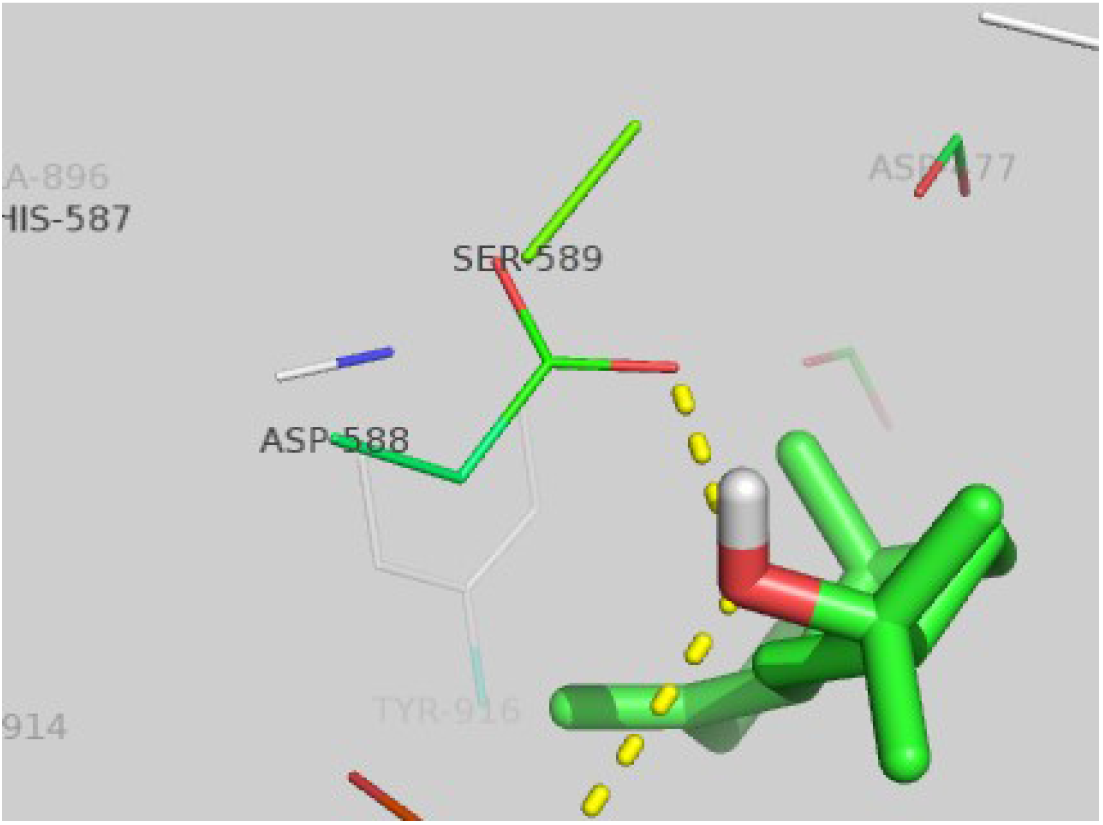
Interaction of Beta Eudesmol.

##### Coulomb energy of Beta eudesmol

The Energy that is associated with the electrostatic forces of the system. The Coulomb energy of Eudesmol was −2258.74 kj/mol. The energy of the protein-ligand complex is stable (Fig 14).

**Figure – 14.**
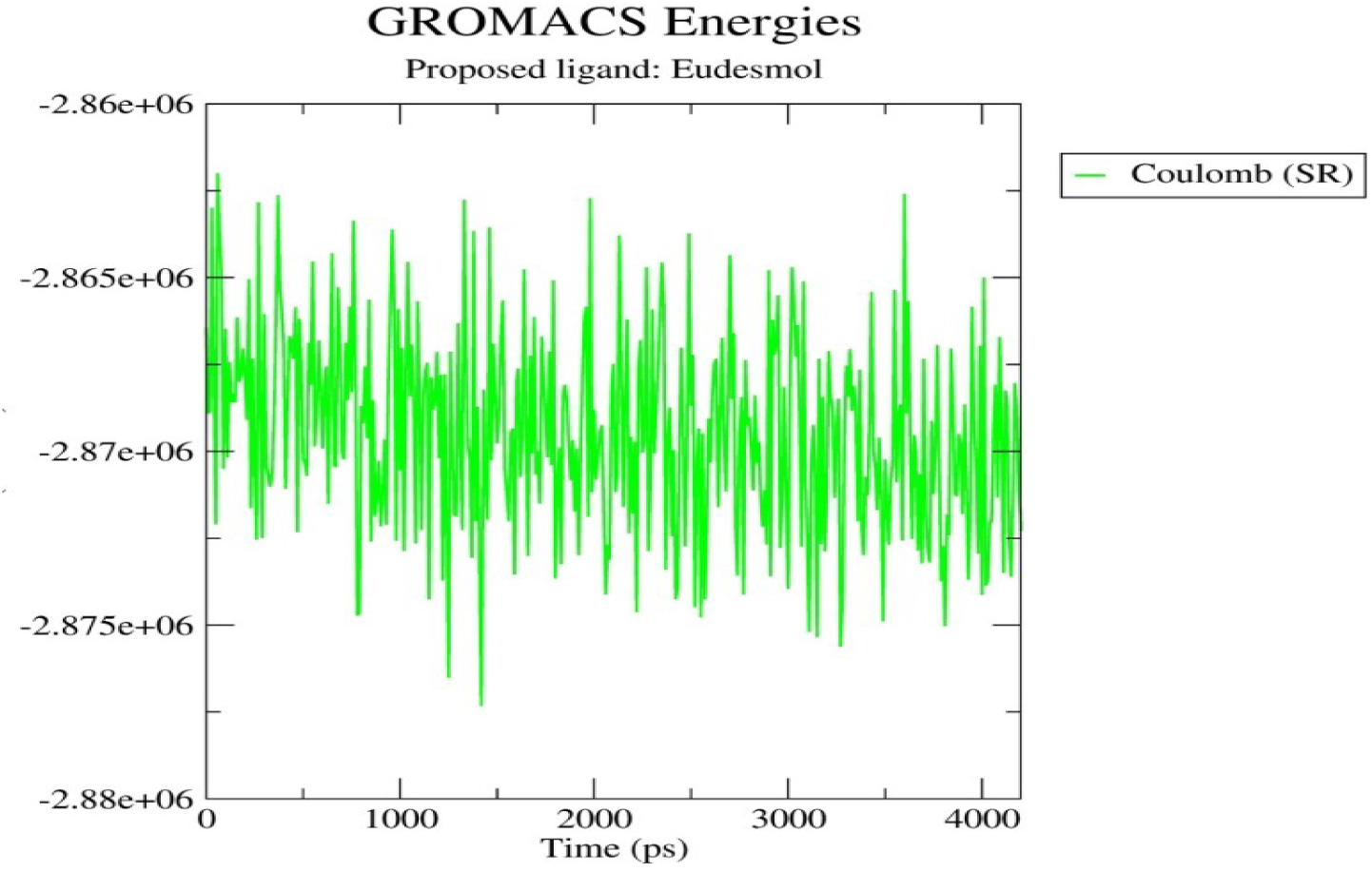
Coulomb energy graph of Beta Eudesmol.

##### RMSD of Beta Eudesmol

The backbone RMSD shows that the equilibrium of the ligand is obtained between 2.5ns to 4ns (Fig 15).

**Figure – 15.**
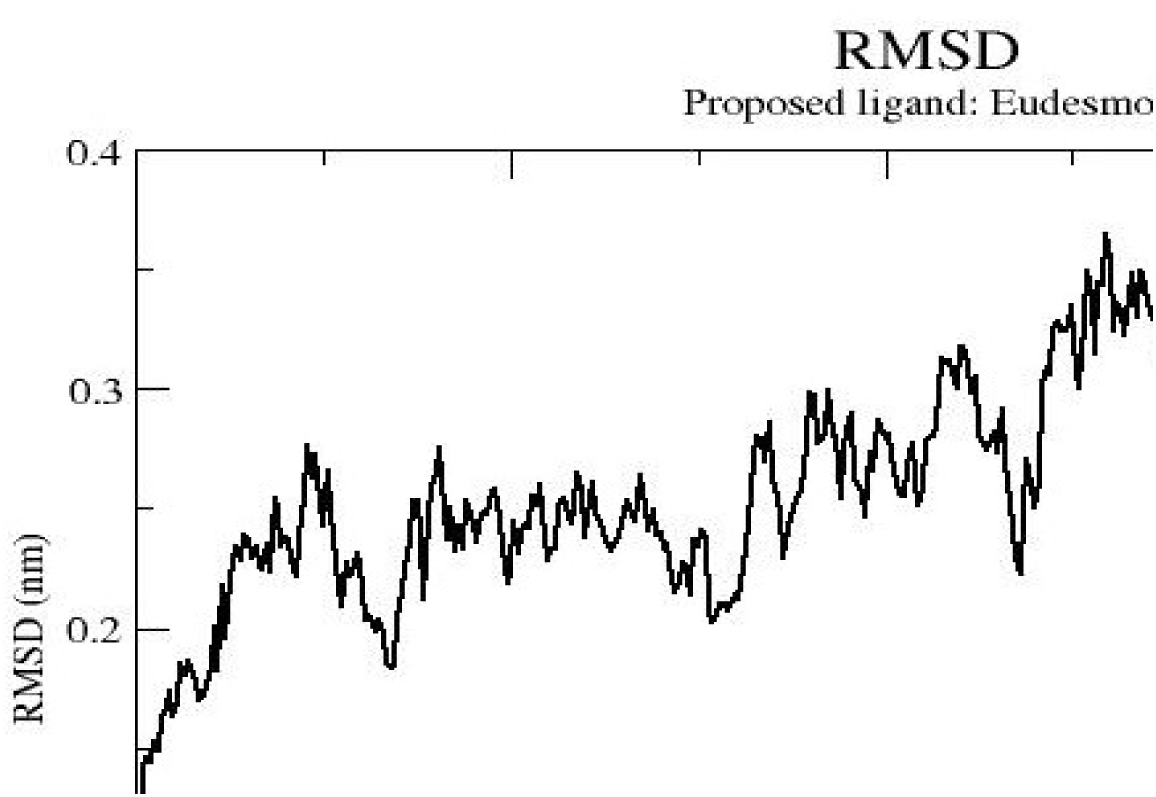
RMSD of Beta Eudesmol.

##### RMSF of Beta Eudesmol

The highly fluctuating residue is **ILU726**. There is no much high fluctuation hence the complex is very stable (Fig 16).

**Figure – 16.**
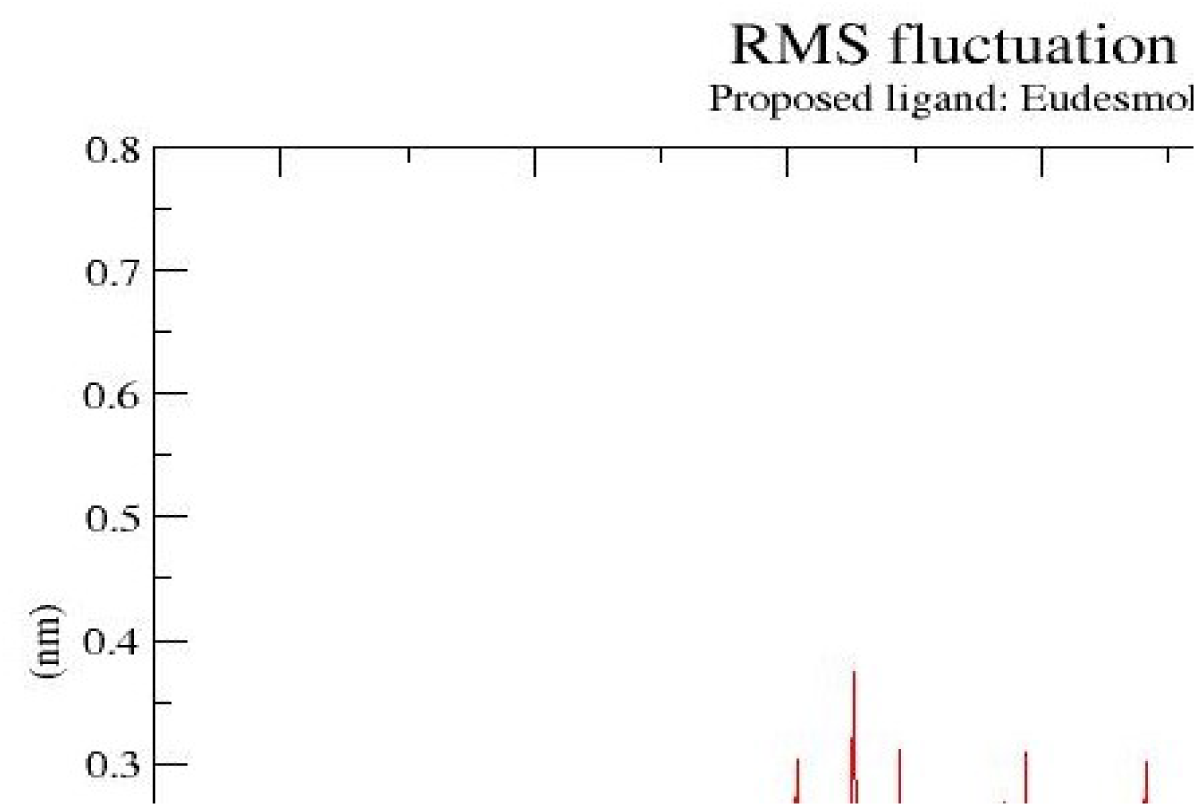
RMSF of Beta Eudesmol.

##### Radius of Gyration of Beta Eudesmol

The compactness of the ligand was obtained from the radius of gyration (Fig 17).

**Figure – 17.**
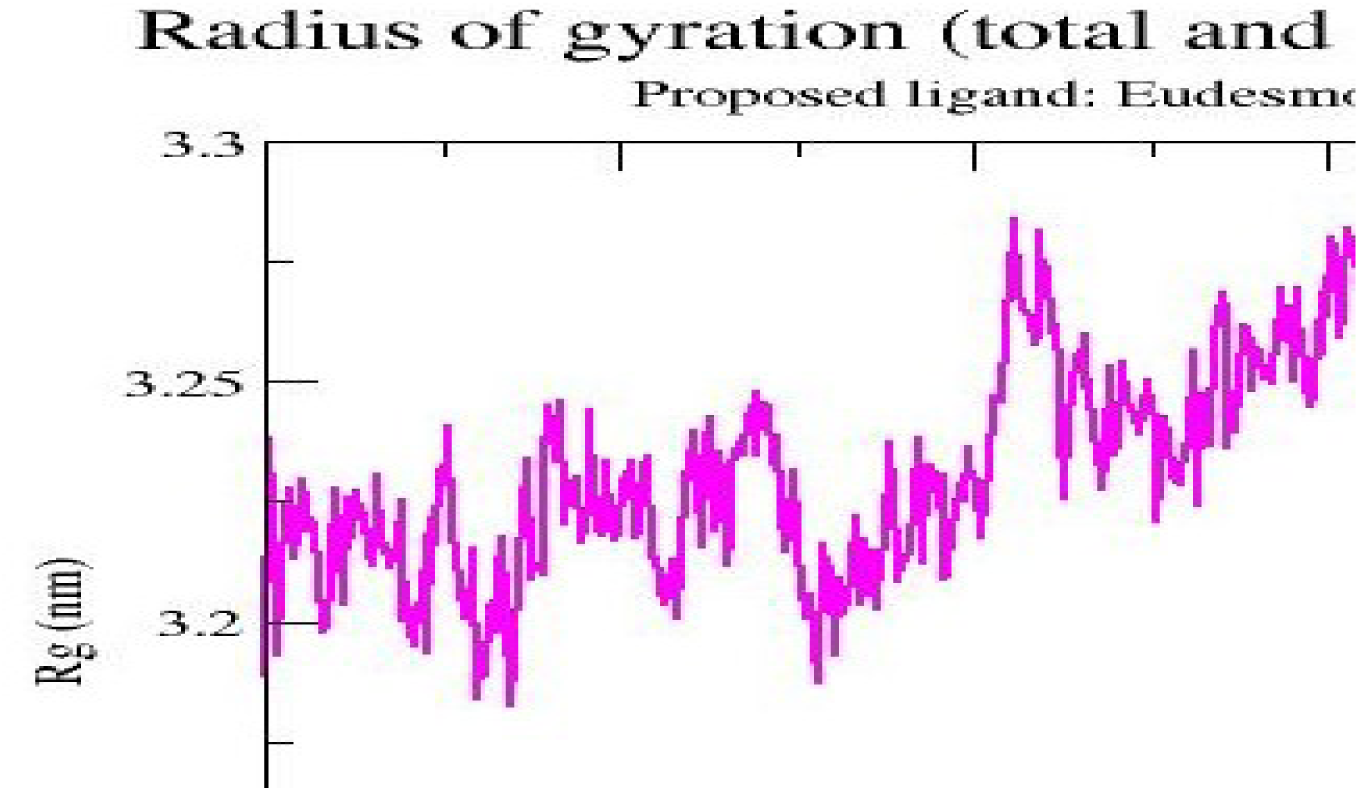
Radius of gyration of Beta eudesmol.

#### 3.2.3. Molecular Dynamic Simulation of Reference Ligand G43 with protein

##### Potential energy of Reference Ligand G43

The potential energy of the reference ligand was obtained as −975.742 kJ/mol. The energy is not that much highly negative as compare to the other proposed ligand (Fig 18). Hence the proposed ligand is more stable than the reference ligand.

**Figure – 18.**
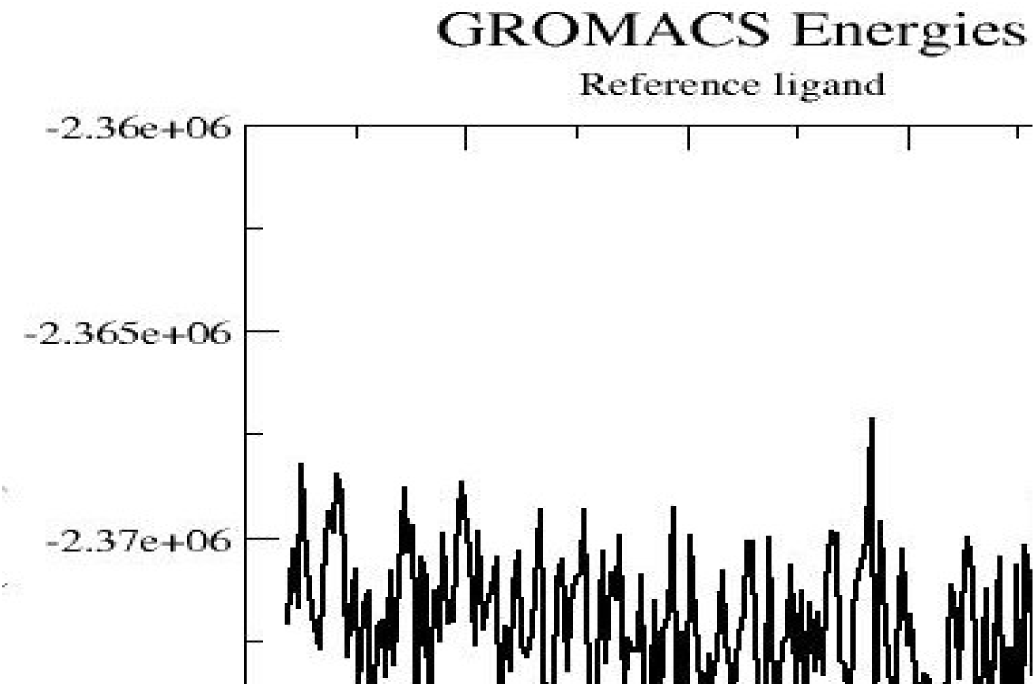
Potential energy of Reference Ligand G43.

**Figure – 19.**
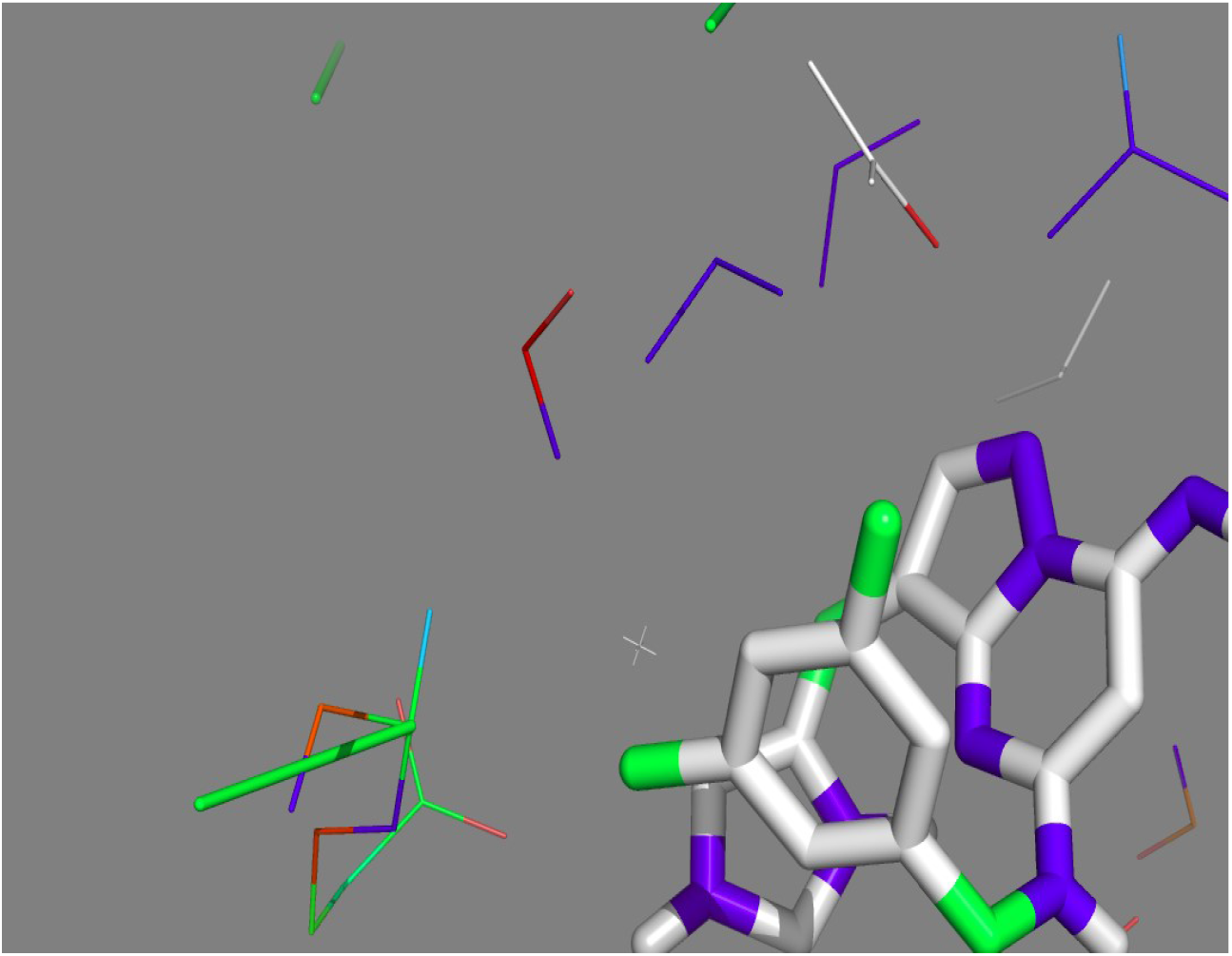
Simulation of Reference Ligand G43 at 0 ns.

##### Coulomb energy of Reference Ligand G43

The coulomb energy was obtained to be −1044.93 kj/mol (Fig 20).

**Figure – 20.**
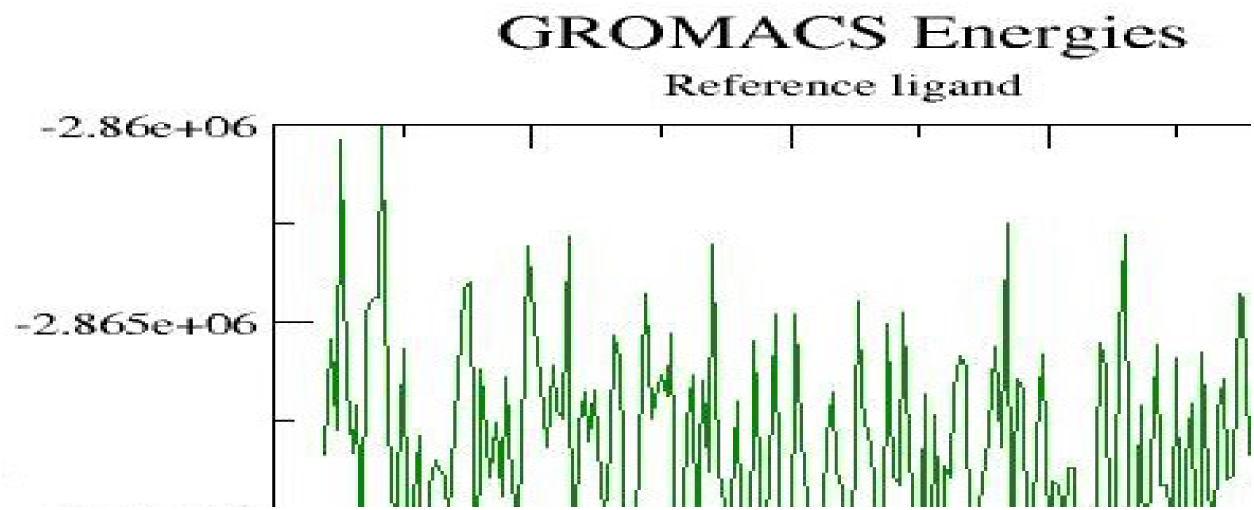
Coulomb energy of Reference Ligand G43.

Lowest potential energy was attained at 2.1 ns. The interacting residues are **ASP477, TYR610**.

**Figure – 21.**
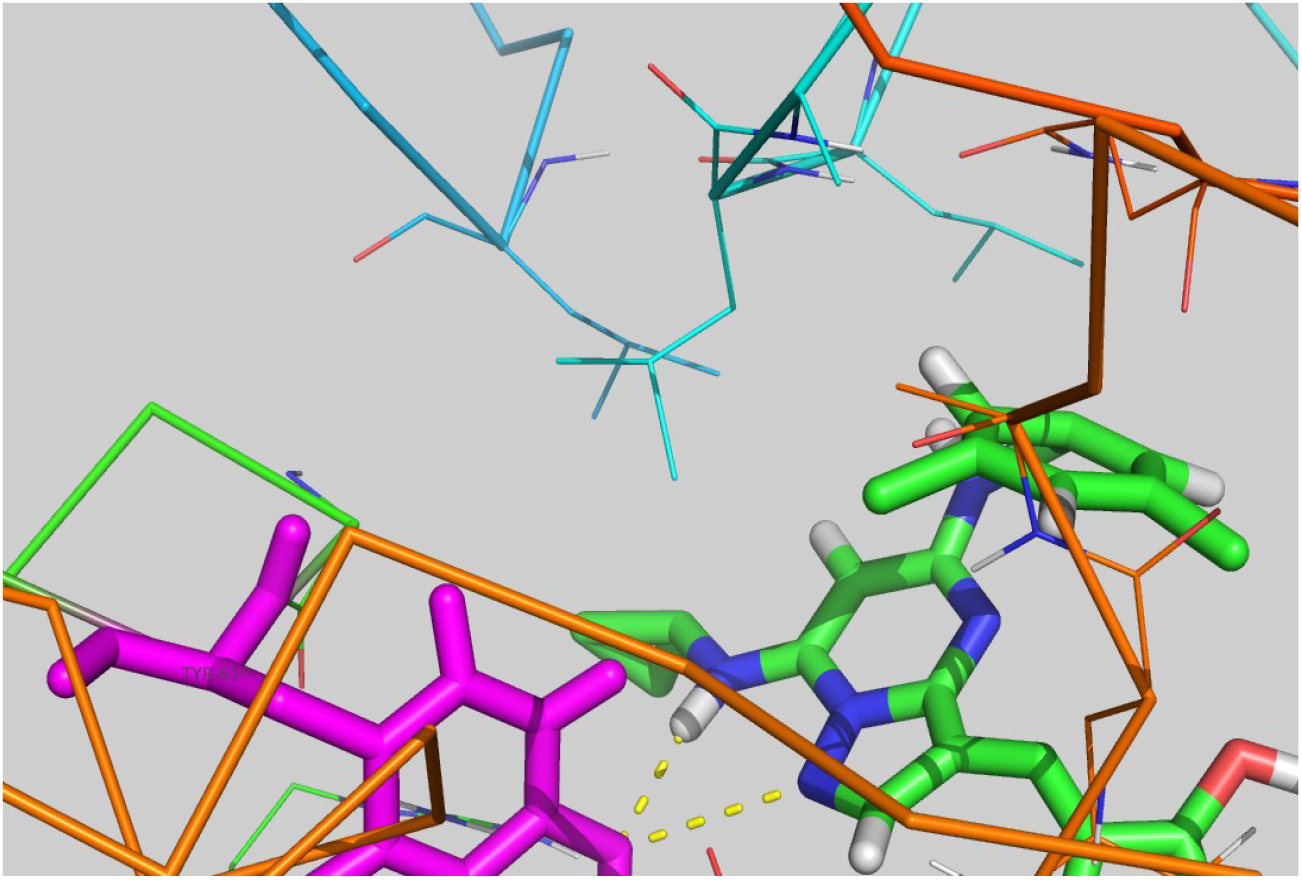
Interacting residues of Reference Ligand G43.

##### RMSD of Reference Ligand G43

The backbone RMSD shows that equilibrium is obtained between 1.7ns to 2.1ns. The stability is obtained and hence the complex backbone is stable (Fig 22).

**Figure – 22.**
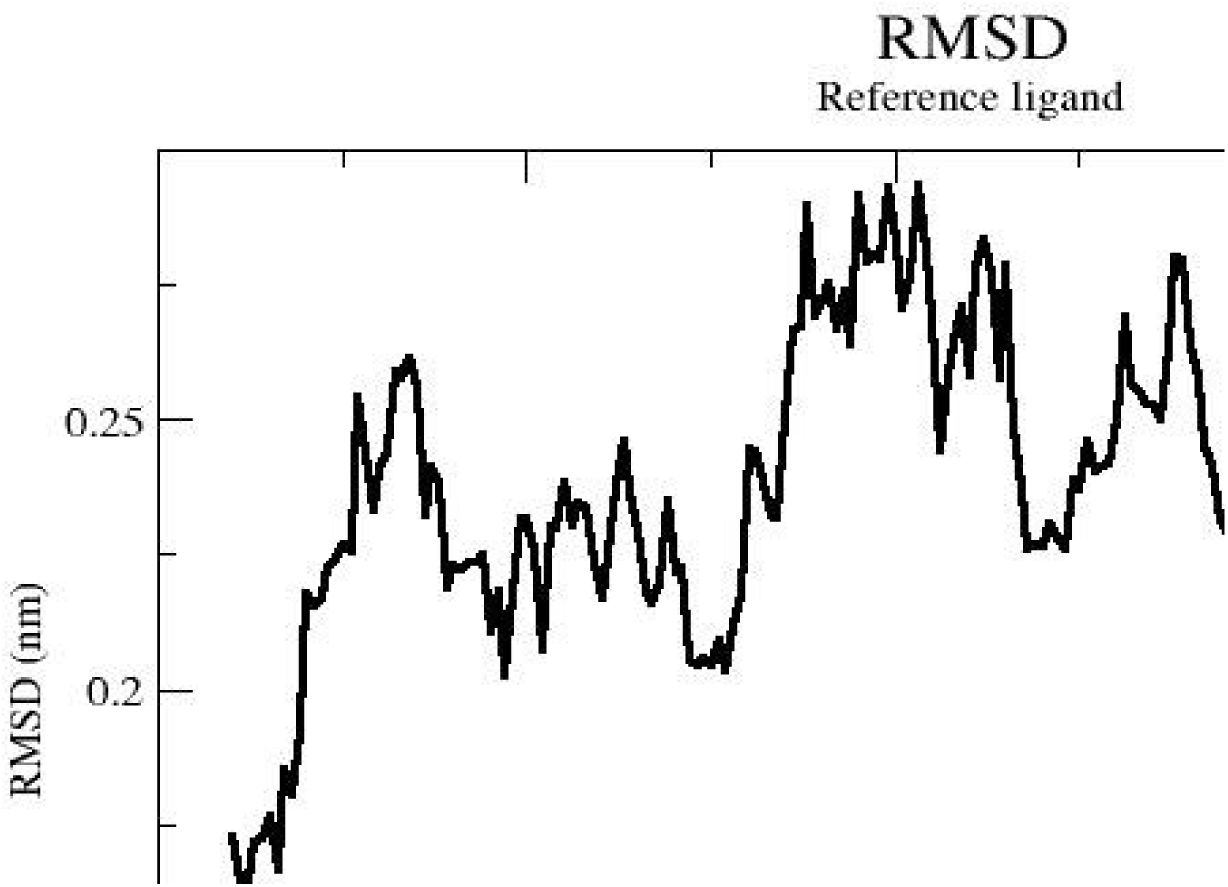
RMSD of Reference Ligand G43.

##### RMSF of Reference Ligand G43

The highly fluctuating residue is **THR425 (Fig 23)**.

**Figure – 23.**
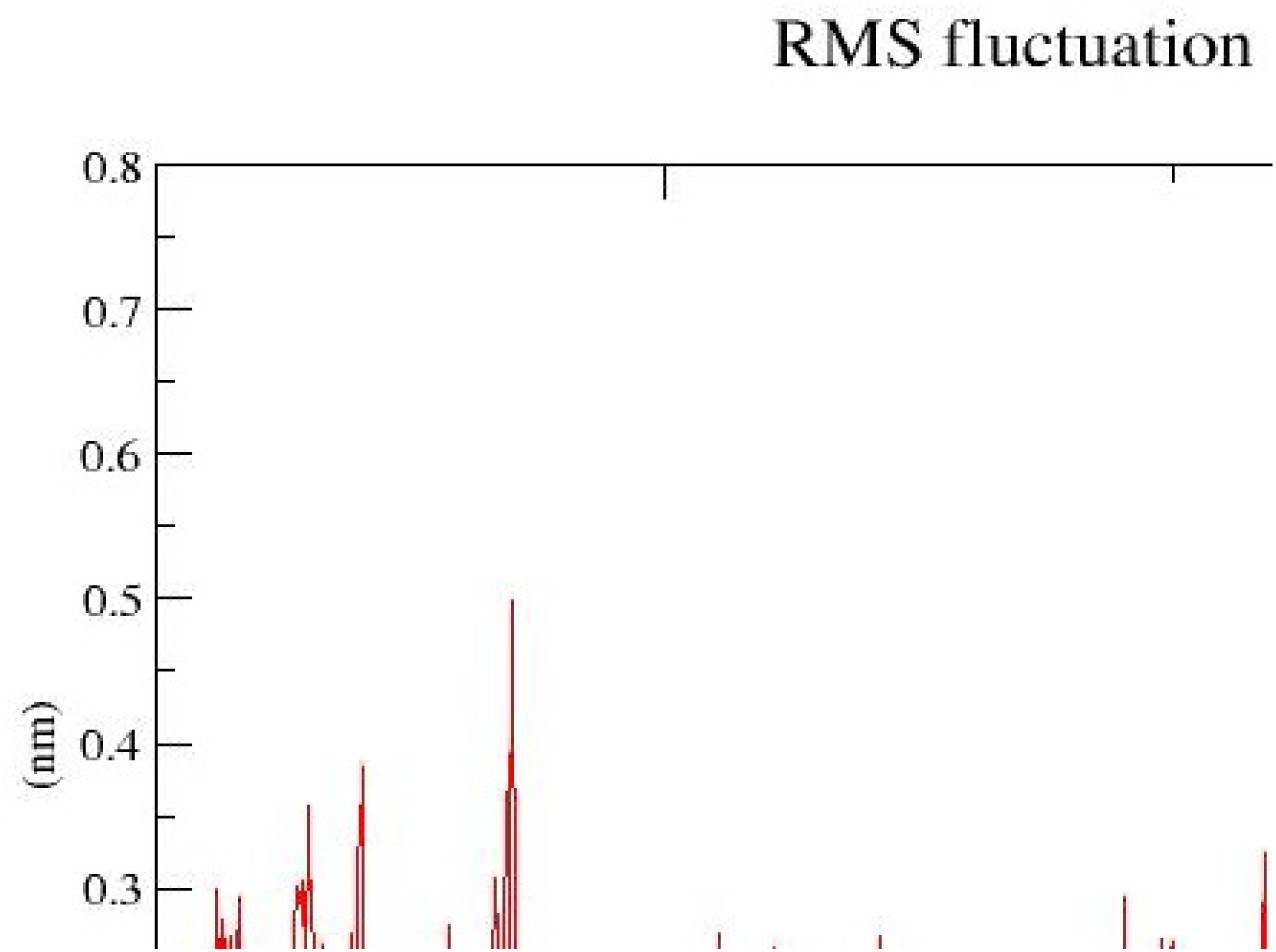
RMSF of Reference Ligand G43.

##### Radius of gyration of Reference Ligand G43

The radius of gyration is shown in Fig 24.

**Figure – 24.**
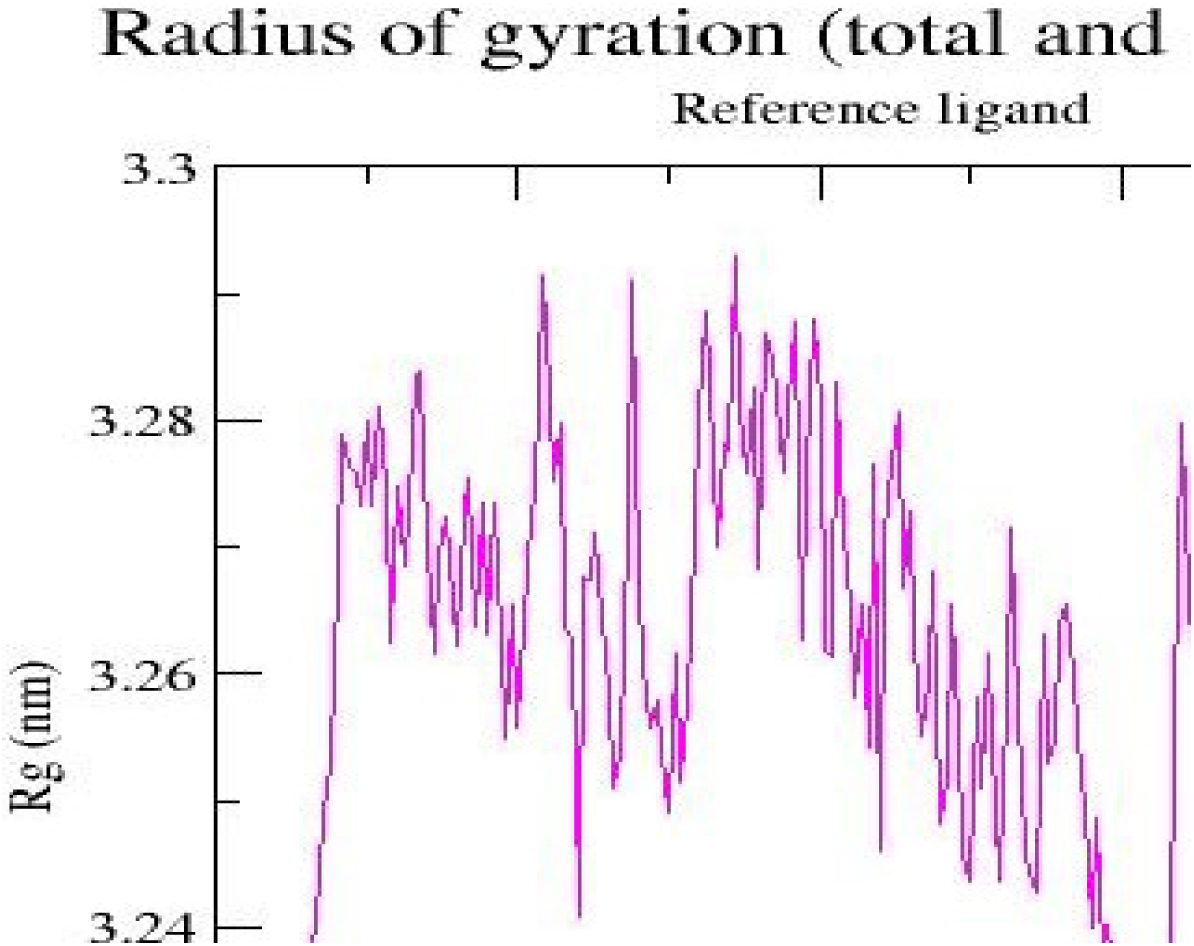
Radius of gyration of Reference Ligand G43.

### 3.3 Classification model of small molecules

Here, the first row molecules are active molecules against glucosyltransferase. In contrast, the second row molecules are inactive against glucosyltransferase (Fig 25).

**Figure – 25.**
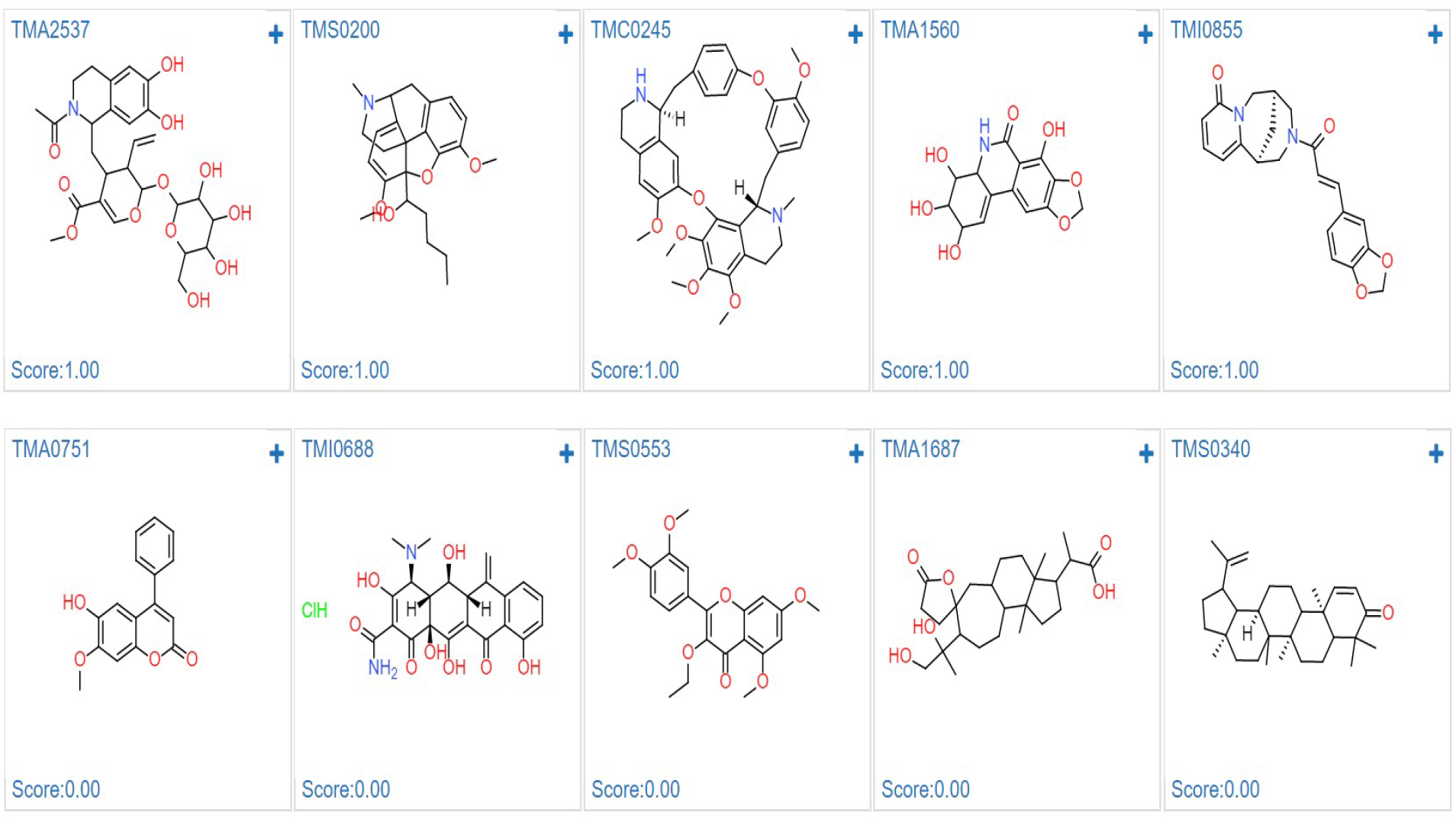
Top 5 Actives (with score 1) & In-actives (with score 0)

#### Accuracy of Classification model

Classification accuracy (or simply, accuracy) is an evaluation metric that is a portion of correct predictions out of the total number of predictions produced by a ML model: Accuracy=Number of correct predictions / Total number of predictions Accuracy can range from 0 to 1 (or 0 to 100%)

**Figure – 26.**
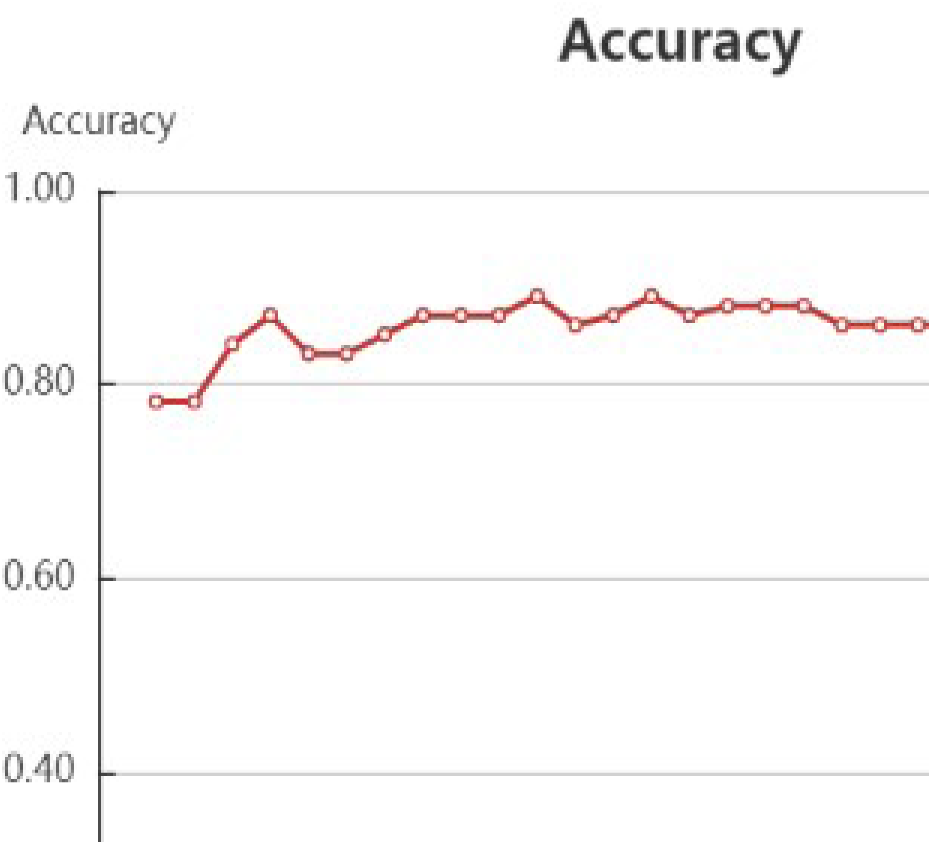
Accuracy of Classification model.

#### Precision of Classification model

Precision is the ratio of true positives to all the positives predicted by the model.

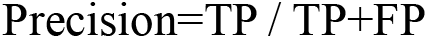

Where, TP is the True Positive & FP is the False Positive.

Low Precision: The more false positives the model predicts, the lower the precision.

**Figure – 27.**
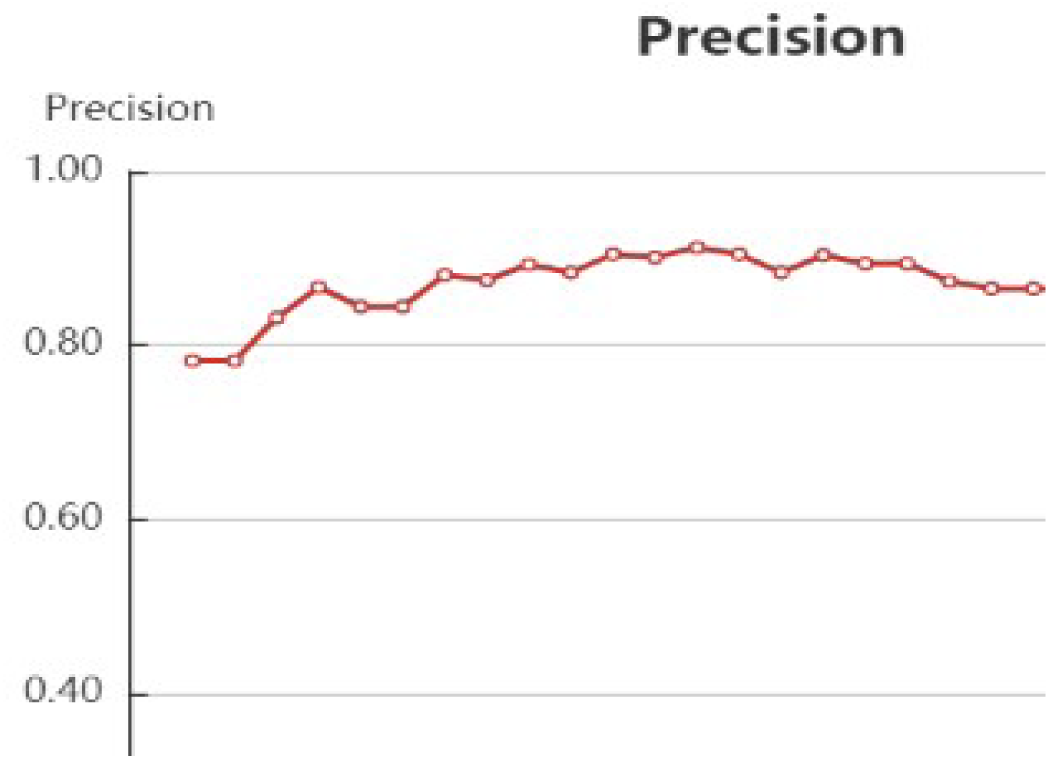
Precision of Classification model.

#### AUC of Classification model

AUC score is the evaluation metric that helps in this case of a single number metric for comparison. AUC stands for **Area Under Curve** and it is simply the percentage of the plot box that lies under the ROC curve. AUC measures the separability. Higher the AUC, better the model would be. The closer it is to 1 (100%), the better

**Figure – 28.**
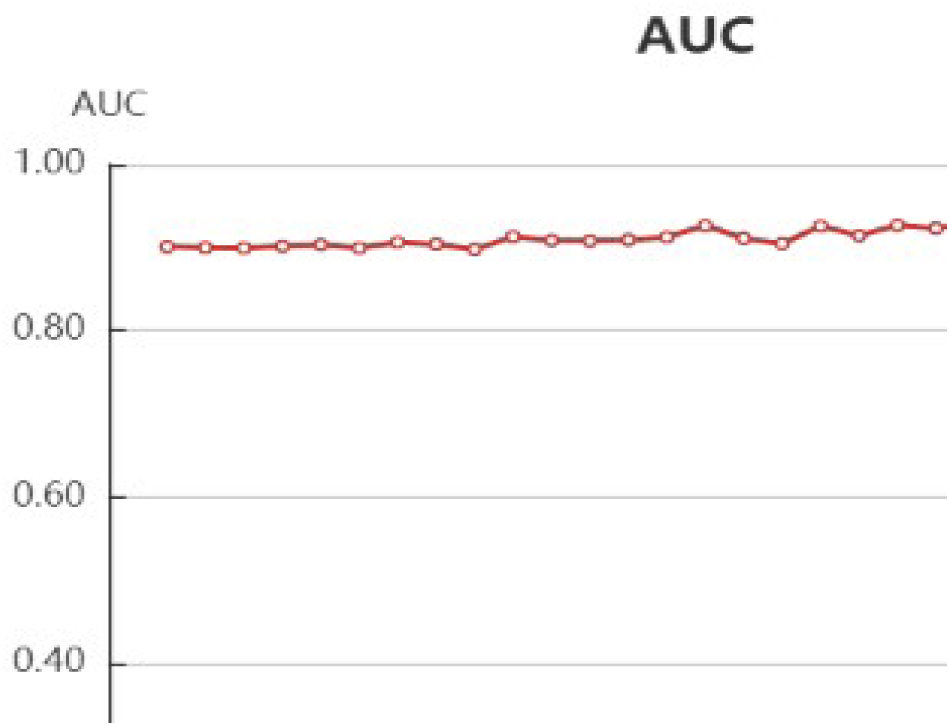
AUC of Classification model.

#### MCC of Classification model

The Matthews correlation coefficient (MCC) is used in machine learning as a measure of the quality of binary (two-class) classifications. The coefficient takes into account true and false positives and negatives and is generally regarded as a balanced measure which can be used even if the classes are of very different sizes. The MCC is in essence a correlation coefficient between the observed and predicted binary classifications; it returns a value between −1 and +1. A coefficient of +1 represents a perfect prediction, 0 no better than random prediction and −1 indicates total disagreement between prediction and observation.

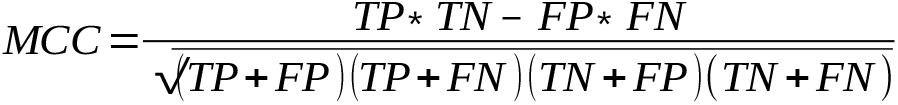

**Figure – 29.**
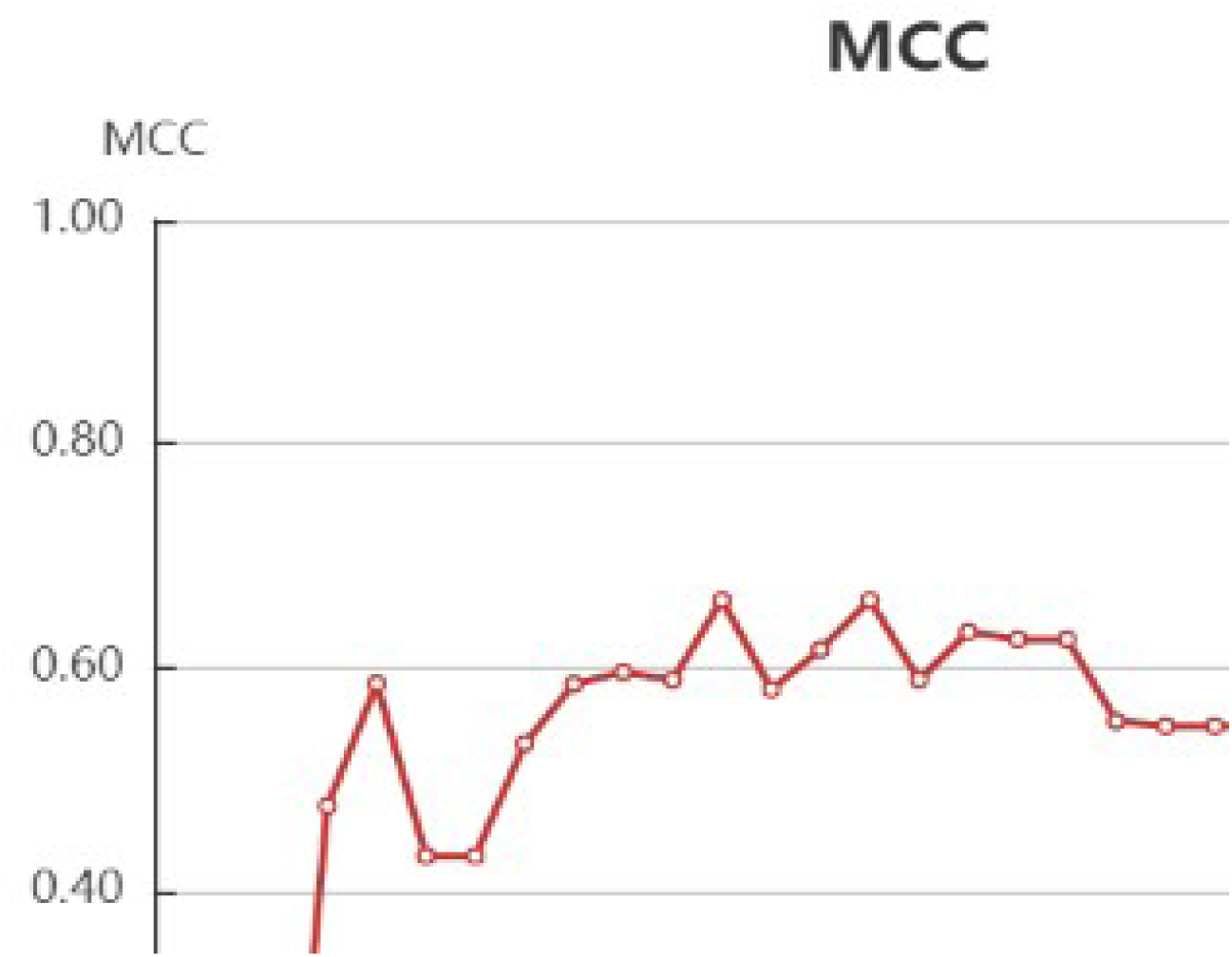
MCC of Classification model.

#### Loss of Classification model

Log Loss, is another evaluation metric that takes into account the output probabilities. It is commonly used as a loss function. Log Loss quantifies the performance of a classifier by penalizing false classifications and taking into account the uncertainty of the predictions which gives a more nuanced view into the performance of the model compared to accuracy. It should be least to refer the model is good.

**Figure – 30.**
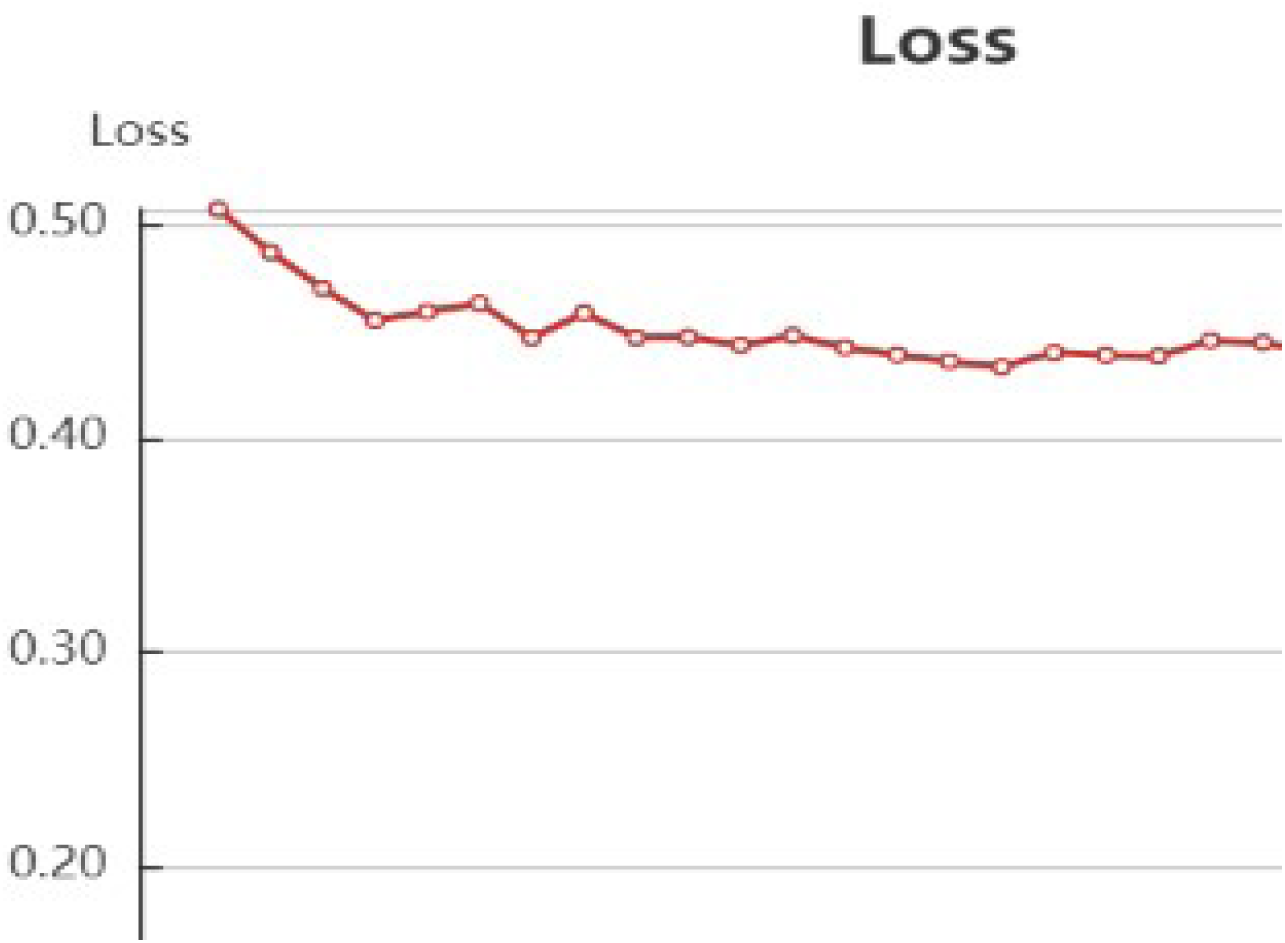
Loss of Classification model.

#### Classification model metrics

At the 30^th^ cycle, the classification model improved with AUC: 0.919, MCC: 0.656, F1: 0.933, precision: 0.895, accuracy: 0.8911 and test loss: 0.422.

| Epochs | Loss | Accuracy | Recall | Precision | F1 | MCC | AUC |
| --- | --- | --- | --- | --- | --- | --- | --- |
| 1 | 0.5065459875 | 0.7822 | 1 | 0.7822 | 0.8778 | 0 | 0.901 |
| 2 | 0.4865402727 | 0.7822 | 1 | 0.7822 | 0.8778 | 0 | 0.8999 |
| 3 | 0.4697061789 | 0.8416 | 1 | 0.8316 | 0.908 | 0.4762 | 0.8993 |
| 4 | 0.454680745 | 0.8713 | 0.9873 | 0.8667 | 0.9231 | 0.5855 | 0.9016 |
| 5 | 0.4588613345 | 0.8317 | 0.962 | 0.8444 | 0.8994 | 0.4315 | 0.9033 |
| 6 | 0.4627904007 | 0.8317 | 0.962 | 0.8444 | 0.8994 | 0.4315 | 0.8993 |
| 7 | 0.446676547 | 0.8515 | 0.9367 | 0.881 | 0.908 | 0.5319 | 0.9062 |
| 8 | 0.4582040086 | 0.8713 | 0.9747 | 0.875 | 0.9222 | 0.5851 | 0.9039 |
| 9 | 0.4467421924 | 0.8713 | 0.9494 | 0.8929 | 0.9202 | 0.596 | 0.8976 |
| 10 | 0.4469831037 | 0.8713 | 0.962 | 0.8837 | 0.9212 | 0.589 | 0.9131 |
| 11 | 0.4433129471 | 0.8911 | 0.962 | 0.9048 | 0.9325 | 0.6602 | 0.9085 |
| 12 | 0.4475990876 | 0.8614 | 0.9241 | 0.9012 | 0.9125 | 0.5805 | 0.9079 |
| 13 | 0.4421242653 | 0.8713 | 0.9241 | 0.9125 | 0.9182 | 0.6162 | 0.9091 |
| 14 | 0.438618431 | 0.8911 | 0.962 | 0.9048 | 0.9325 | 0.6602 | 0.9125 |
| 15 | 0.4356696936 | 0.8713 | 0.962 | 0.8837 | 0.9212 | 0.589 | 0.9264 |
| 16 | 0.4331509949 | 0.8812 | 0.9494 | 0.9036 | 0.9259 | 0.6318 | 0.9108 |
| 17 | 0.4398030621 | 0.8812 | 0.962 | 0.8941 | 0.9268 | 0.6251 | 0.9045 |
| 18 | 0.4384767171 | 0.8812 | 0.962 | 0.8941 | 0.9268 | 0.6251 | 0.9258 |
| 19 | 0.4380863008 | 0.8614 | 0.962 | 0.8736 | 0.9157 | 0.5519 | 0.9143 |
| 20 | 0.4453485354 | 0.8614 | 0.9747 | 0.8652 | 0.9167 | 0.5476 | 0.9269 |
| 21 | 0.4444094141 | 0.8614 | 0.9747 | 0.8652 | 0.9167 | 0.5476 | 0.9229 |
| 22 | 0.4396597631 | 0.8614 | 0.9747 | 0.8652 | 0.9167 | 0.5476 | 0.931 |
| 23 | 0.4245092869 | 0.8911 | 0.9747 | 0.8953 | 0.9333 | 0.6565 | 0.9177 |
| 24 | 0.4276346889 | 0.8812 | 0.962 | 0.8941 | 0.9268 | 0.6251 | 0.9275 |
| 25 | 0.4340768864 | 0.8713 | 0.9747 | 0.875 | 0.9222 | 0.5851 | 0.9266 |
| 26 | 0.4224902262 | 0.8812 | 0.962 | 0.8941 | 0.9268 | 0.6251 | 0.9278 |
| 27 | 0.4188840531 | 0.8911 | 0.962 | 0.9048 | 0.9325 | 0.6602 | 0.9189 |
| 28 | 0.4215655858 | 0.8911 | 0.9747 | 0.8953 | 0.9333 | 0.6565 | 0.918 |
| 29 | 0.4198817853 | 0.8911 | 0.9747 | 0.8953 | 0.9333 | 0.6565 | 0.9171 |
| 30 | 0.4229586337 | 0.8911 | 0.9747 | 0.8953 | 0.9333 | 0.6565 | 0.9194 |

## 4. CONCLUSION

The toxicity of Eudesmol was predicted to be low by Tox tree tool and G43 was predicted to be highly toxic. The potential energy of Glycosyl transferase-Eudesmol (−1500 kJ/mol) indicates its higher stability as compared to Glycosyl tranferase-G43 complex (−1100kj/mol). The RMSD, RMSF and Radius of gyration have been plotted for Glycosyl transferase, Glycosyl transferase-Eudesmol and Glycosyl tranferase-G43 complex. Comparatively, Glycosyl transferase-Eudesmol shows more stability during 4.2 ns. The average interaction energy between Glycosyl transferase - Eudesmol was −120kj/mol. The set of active and inactive molecules were taken from deep learning server and top 5 actives and inactive molecules were predicted for the enzyme Glycosyl transferase.

## Acknowledgements

We acknowledge SASTRA Deemed University for providing computational facility.

